# The investigation of the role of VHL-HIF signaling in DNA repair and apoptosis in zebrafish

**DOI:** 10.1101/767459

**Authors:** HR Kim, K Santhakumar, E Markham, D Baldera, HE Bryant, SF El-Khamisy, FJ van Eeden

**Affiliations:** Bateson Centre, Firth Court, University of Sheffield, Western Bank, Sheffield S10 2TN, UK; Department of Genetic Engineering, SRM Institute of Science and Technology, Kattankulathur-603203, India; Department of Oncology & Metabolism, The Medical School, Beech Hill Road, Sheffield S10 2RX, UK; Department of Molecular biology and Biotechnology, Firth Court, University of Sheffield, Western Bank, Sheffield S10 2TN, UK

**Keywords:** Hif, Vhl, DNA repair, apoptosis, chemo/radio-resistance

## Abstract

pVHL is a tumor suppressor. The lack of its function leads to various tumors, among which ccRCC (clear cell renal cell carcinoma) has the most serious outcome due to its resistance to chemotherapies and radiotherapies. Although HIF promotes the progression of ccRCC, the precise mechanism by which the loss of VHL leads to tumor initiation remains unclear. We exploited two zebrafish *vhl* mutants, *vhl* and *vll*, and *Tg(phd3::GFP)^i144^* fish to identify crucial functions of Vhl in tumor initiation. Through the mutant analysis, we found that the role of pVHL in DNA repair is conserved in zebrafish Vll. Interestingly, we also discovered that Hif activation strongly suppressed genotoxic stress induced DNA repair defects and apoptosis in *vll* and *brca2* mutants and in the embryos lacking ATM activity. These results suggest the potential of HIF as a clinical modulator that can protect cells from accumulating DNA damage and apoptosis which can lead to cancers and neurodegenerative disorders.

## Introduction

In humans, mutations in the pVHL protein, a tumor suppressor protein, predispose patients to Von Hippel Lindau (VHL) disease, a rare form of dominantly inherited cancer syndrome. The patients suffer from recurrent cysts and tumors that can develop into malignant tumors in multiple tissues and organs throughout their lifetime. These include hemangioblastomas of retina and the central nervous system, cysts in pancreas and kidneys, pheochromocytoma and clear cell renal cell carcinoma (ccRCC). Among these, ccRCC is the leading cause of death from the VHL disease. It is difficult to be diagnosed at early stages when the tumors are localized and can be treated by surgery, and the advanced forms of tumor are extremely resistant to chemotherapies and radiation therapies. pVHL is also mutated in a majority of cases of the sporadic form of ccRCC, indicating that VHL is a major tumor driver in ccRCC [1, 2].

pVHL is best known as a negative regulator of Hypoxia Inducible Factor (HIF) transcription factors that play crucial roles for the adaptation of cells to the hypoxic environment. HIF transcription factors consist of two subunits, oxygen sensitive α and constitutively present β. In a normal oxygen tension environment, HIFα is hydroxylated by oxygen sensing enzymes, Prolyl Hydroxylases (PHDs) that use oxygen as substrates. Once it is hydroxylated, HIFα is constantly degraded by E3 ubiquitin ligase complex where pVHL is a substrate recognition subunit. However when cells are exposed to hypoxia, the inactivation of PHD enzymes lead to HIFα stabilization, its translocation into the nuclei and the formation of functional heterodimers with HIF1β subunit. HIF heterodimers bind to the HIF responsive element (HRE) in their target genes to activate genes that are involved in the adaptation process in hypoxia such as angiogenesis, erythropoiesis, glucose uptake, glycolysis, cell proliferation and cell survival [3, 4].

In the absence of VHL, regardless of the environmental oxygen tension, HIF is constantly activated mimicking hypoxic conditions. Tumors are often hypoxic at their core and the activated HIF is known to be advantageous for the survival of tumor cells and to be associated with poor prognosis in various cancers including ccRCC [5]. Understanding the molecular pathways of ccRCC led to the development of a few target therapies. Therapies that target angiogenesis have been a logical approach to treat ccRCC, but only have temporary effect that are accompanied by significant side effects. Alternative approaches may be developed through deeper understanding of the role of VHL.

There are a multitude of HIF independent functions of pVHL that could be potentially targeted for ccRCC treatment, although these are not as intensively investigated as HIF regulation. These include microtubule stabilization and maintenance of the primary cilium, regulation of the extracellular matrix formation, and control of cell senescence and apoptosis, transcriptional regulation [6] and DNA repair [7, 8]. The mutations in pVHL that cause pheochromocytoma retain the ability to regulate HIF degradation, indicating that the HIF independent role of pVHL is important for tumorigenesis [9]. Also, activated HIF alone cannot initiate kidney cancer development in humans and animal models, suggesting that HIF independent roles of pVHL could be important for the initiation of ccRCC development [10, 11].

Besides accumulated HIF, genomic instability is one of characteristics of ccRCC. Evidence suggests that the accumulation of DNA lesions in our body is associated with cancer development and aging [12]. Indeed, inherited mutations in the genes that play important roles in the DNA repair pathways predispose patients to cancer development or premature aging [13]. For example, mutations in one allele of the *BRCA2* gene that promotes double strand break repair (DSB) by homologous recombination (HR), increase susceptibility to breast and ovarian cancer, and mutations in ATM that is crucial for DNA repair and cell cycle control upon DNA damage, cause Ataxia -Telangiectasia (A-T) syndrome that is characterized by the very high risk of malignancy, radiosensitivity and progressive ataxia. Heterozygous individuals have an increased risk of cancer [14, 15]. In line with this, Metcalf *et al*. demonstrated that pVHL is required for DSB repair in ccRCC cell lines, implicating the role of pVHL in DNA damage repair as a cause of the genomic instability in ccRCC [7]. In this study, pVHL was hypothesized to work together with ATM upon DNA damage to trigger downstream signaling for the recruitment of DNA repair proteins at the site of DNA damage.

Similarly, Scanlon *et al*. demonstrated that there was an accumulation of DNA damage in *VHL* defective ccRCC cell lines compared to *VHL* complemented cells [8]. There was also downregulation of genes that regulate DSB repair and mismatch repair (MMR) in the ccRCC cells, that may explain the increase in the DNA damage seen. The authors suggest that the VHL deficient cells activate processes that are similar to those in the cells exposed to hypoxia. It was speculated that the downregulation of DNA repair genes in ccRCC cell lines is due to the activation of HIF2 rather than HIF1, since ccRCC cells expressing only HIF2 exhibit the same gene expression profile, as that of the cells expressing both HIF transcription factors, i.e. downregulated DNA repair genes. This study also demonstrated the increased sensitivity of ccRCC cells to PARP inhibitor likely because of the DSBR defect in the ccRCC cells.

Therefore, it is clear that the role of VHL in the DNA repair is associated with ccRCC development. However, there are fundamental discrepancies in the above two studies. Although both studies were performed using ccRCC cell lines, Metcalfe *et al*. reported that the role of VHL in DSBR is HIF independent, whereas Scanlon *et al*. suggested that the defects in the DNA repair in the *vhl* mutant cells are similar to those in the cells exposed to hypoxia and they are likely to involve HIF2 transcription factor. This is the limitation of the studies using isolated cells: the cells accumulate mutations in the adaptation process and become different from the tumours they are originated from, although cell lines have certainly been extremely valuable in identifying cancer drugs (*e.g*., PARP inhibitors) [16]. In addition, the drug responses in the cell lines often cannot be recapitulated in human clinical trials [17, 18]. Therefore, to consolidate the findings in the cell lines we need a whole organism model in which cells maintain their contextual environment and do not undergo the adaptation process in the laboratory environment. Besides, already transformed cancer cell lines are difficult to use for studying the cancer initiation.

More importantly, ccRCC is notorious for its resistance to chemotherapeutic reagents; therefore, it can be speculated that if there is a defect in the DNA repair in the ccRCC as the above two papers suggest, the tumor cells will be extremely sensitive to, rather than resistant to, the DNA damaging reagents further emphasizing the limits of cell lines for studies. We hypothesize that even if the DNA repair function of VHL is important for the initiation of ccRCC development, there must be mechanisms that compromise the effect of the loss of its function and provide resistance to therapies as ccRCC progresses. Indeed, the activation of HIF1α is known to be associated with the resistance of various tumors to chemo and radio therapies by mechanisms that are extremely diverse and complex [19] for instance. Similarly, HIF1α was shown to provide the radioresistence in hypoxic mice mesenchymal stromal cells by upregulating DNA repair proteins [20]. pVHL is also known to regulate p53 that is another crucial transcription factor in the adaptation of cells in response to genotoxic stress and its malfunction provides various tumours with resistance to chemo and radio therapies [21]. Therefore, using zebrafish as a whole organismal model, we aim to understand the role of HIF dependent and independent role of VHL in DNA repair and apoptosis and the role of VHL/HIF in the p53 regulation in response to genotoxic stress.

Zebrafish provides an excellent high throughput vertebrate model system. Nearly 70% of human genes have orthologous genes in zebrafish and when only disease related genes are considered, around 82% of genes are associated with at least one zebrafish orthologue [22]. Zebrafish also provides advantages over higher vertebrate models such as fecundity, *in vitro* fertilisation and easy genetic manipulation. Due to a genome duplication event, there are two zebrafish *vhl* homologues, *vhl* and *vll* (*vhl like*). Previously we reported the role of *vhl* in the HIF regulation and the null zebrafish *vhl* mutant mimics Chuvash polycythemia in human [23–25]. In this report, we generated a mutant for the other *vhl* homologous gene, *vll*, by zinc finger endonuclease and investigated the function of *vhl* and *vll* in the DNA repair. We took advantage of *T*g(*phd3::GFP)^i144^* reporter line which expresses a high level of GFP in the absence of functional *vhl* but a minimal level of GFP in the presence of one wild type allele of *vhl* [26]. We used *vhl+/-; T*g(*phd3::GFP)^i144^* fish as a unique tool to study *in vivo* genomic instability using the *vhl* gene as a “sentinel”, since cells express a high level of GFP when the remaining wild type *vhl* is lost.

Interestingly the function of human VHL in HIF regulation and DNA repair seems to be partially segregated into zebrafish Vhl and Vll respectively, Hif regulation in Vhl and DNA repair in Vll. We found that the role of Vll in the DNA repair is Hif independent. Surprisingly however, we identified a role of Hif in the promotion of DNA repair and protection of embryos from apoptosis when embryos were exposed to genotoxic stress. Upregulated Hif, suppresses not only the DNA repair defects in the *vll* mutants but also the defects in the *brca2* mutants and the embryos in which ATM function is inhibited. We hope our results will provide a better understanding of ccRCC development and open up great possibilities for the HIF activators to be exploited for the treatment of disorders associated with DNA repair defects. We think our *vhl;vll* double mutants can also provide an especially useful system for drug discovery to identify drugs that could overcome the resistance of tumour cells to chemo- and radio therapies.

## Results

### 1. Genetic knock out of *vhl* paralogue *vll*, by target specific endonuclease

There are two zebrafish *vhl* homologues, *vhl* and *vll* (*vhl like*). The function of zebrafish *vhl* has been well characterized and documented [23, 24]. The function of its homologue *vll*, however, has not been reported yet. Therefore, to fully appreciate the function of Vhl in zebrafish, we created a genetic knock-out of *vll* by a zinc finger nuclease. The mutant allele lacks 23 nucleotides around transcription starting site including ATG and it is expected to be functionally null (Fig. 1A). Sequence analysis revealed that zebrafish Vhl shares 70% of homology with human VHL whereas zebrafish Vll shares 52% of homology (Fig. 1B). As we previously reported, *vhl* mutant embryos can survive up to 9 dpf, and the mutant embryos show typical hypoxic responses such as increased blood vessel formation, upregulated erythropoiesis and hyperventilation [23]. On the contrary, *vll* mutant embryos are fertile and fully viable, and show no morphological signs of Hif upregulation (Fig. 1C).

**Fig.1.**
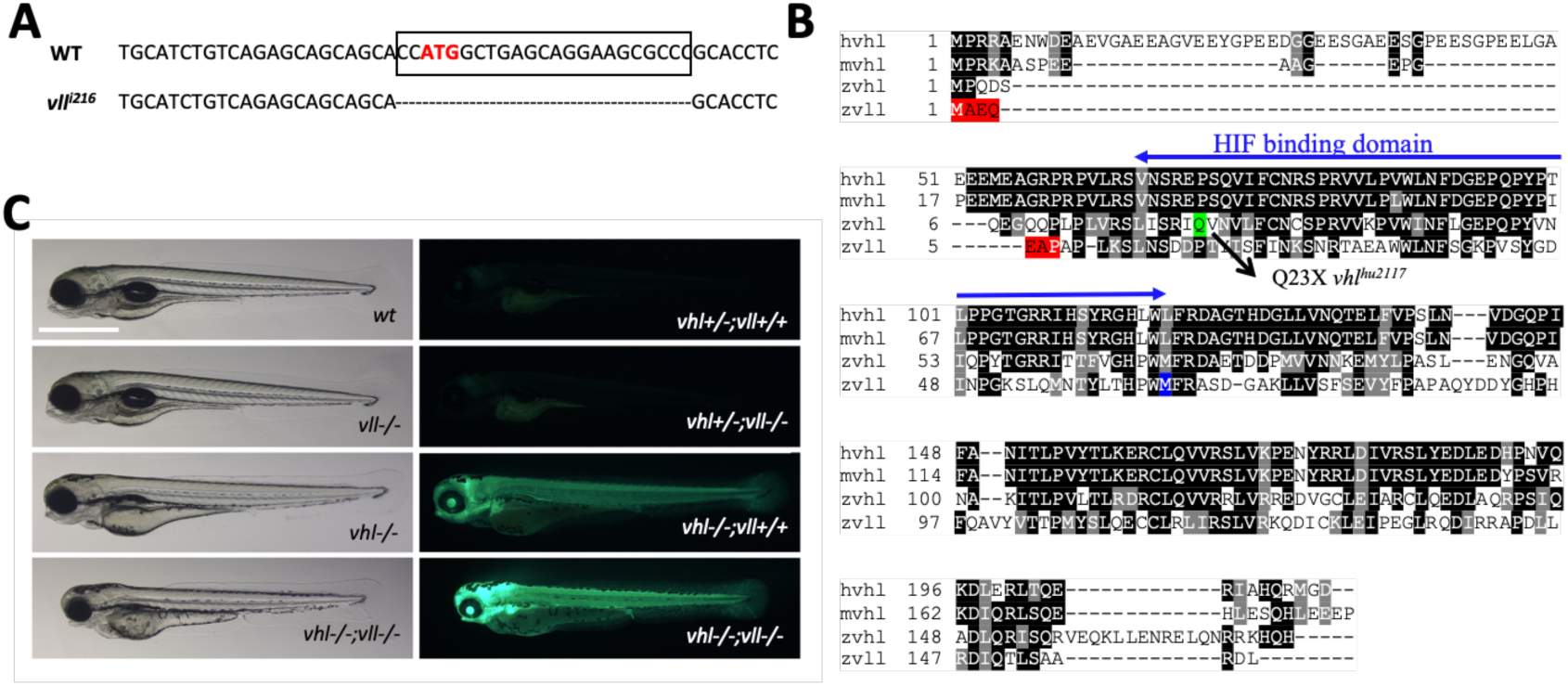
A *vll* mutant allele, *vll^216^*, is generated by a zinc finger nuclease. A. The mutant allele harbors 23 nucleotide deletion around translational starting site. B. Zebrafish Vll shares 52% homology with human VHL. The seven amino acids that are lost in *vll^216^*are highlighted in red. Even if the next available ATG (highlighted in blue) were to be used as an alternative translational start site, it would lead to a protein that lacks most of the predicted Hif binding domain. C. Since *phd3* (*egln3*) expression most dramatically reflects the increased Hif expression, we used *Tg(phd3::EGFP)^i144^* transgenic line as a Hif signaling readout. Zebrafish *vll-/-* embryos do not show any morphological phenotype and the *Tg(phd3::EGFP^)i144^* expression in the mutants was identical to that in the wild type embryos. Reflecting the role of Vll in Hif regulation in the absence of Vhl, there was an enhanced GFP expression in the *vhl-/-;vll-/-* double mutant embryos in comparison to that in *vhl-/-* embryos. Scale bar: 1mm

When we examined the expression of key Hif target genes by qPCR in the *vll*-/- mutant embryos, in spite of the lack of visible phenotypes, there was a modest level of upregulated Hif target genes in the *vll*-/- mutants. Furthermore, there was a remarkable and more than additive upregulation of HIF target genes in the *vhl-/-;vll-/-* double mutants (Table 1). The characteristics of *vhl* mutants by Hif upregulation, such as the reduced usage of yolk and slowdown in growth, were also more enhanced in *vhl-/-;vll-/-* double mutants in comparison to *vhl*-/- single mutants (Fig. 1C). We concluded that, although *vll* is not required for Hif regulation in the presence of *vhl*, it plays an important role in Hif regulation in the absence of *vhl,* sharing a functional redundancy with *vhl*.

**Table 1.**
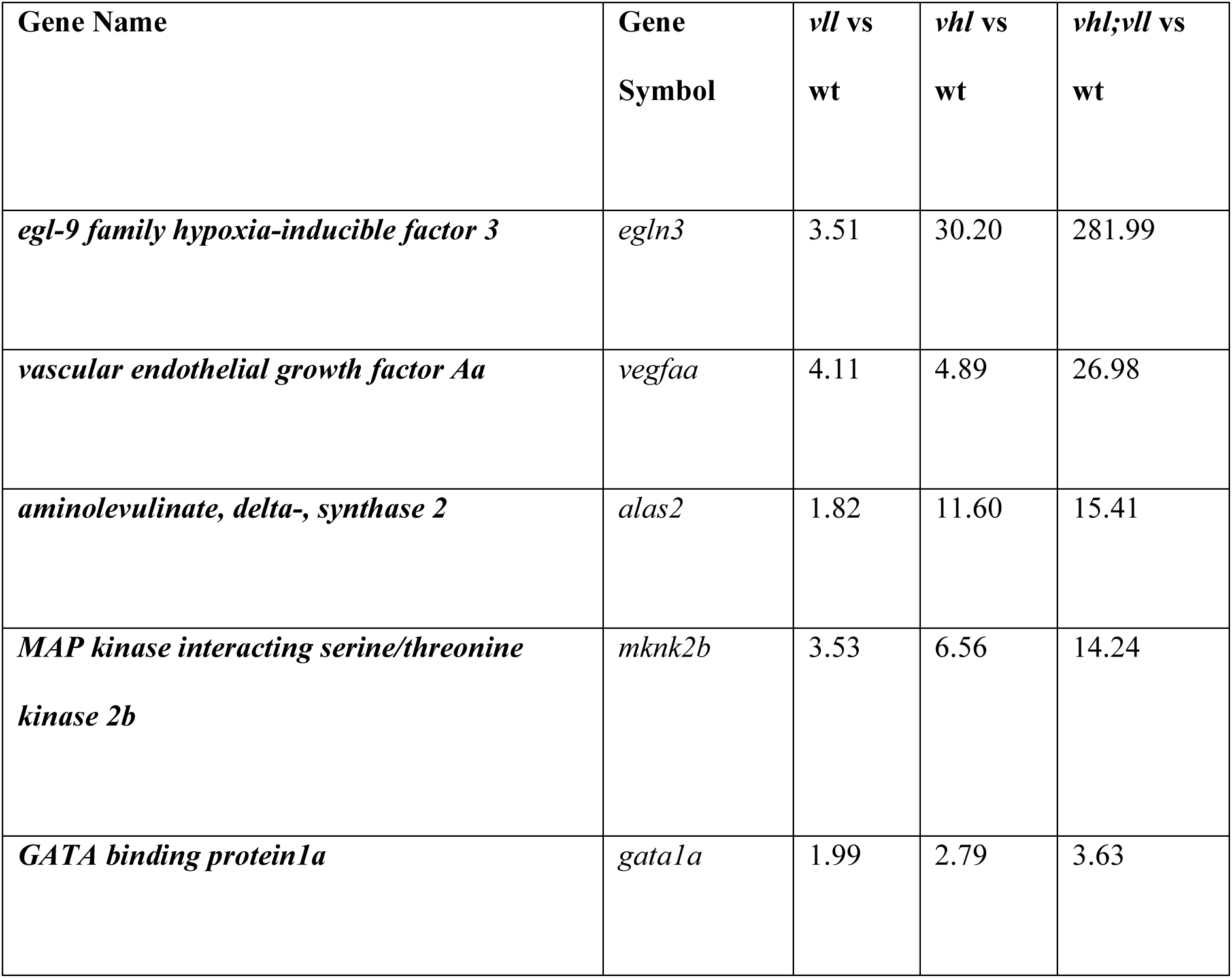
The qPCR results reveal the modest role of Vll and the synergistic effect of the loss of Vhl and Vll in the Hif regulation. We performed qPCR to examine the expression level of a few Hif target genes. There was a modest upregulation of Hif target genes in *vll* mutants indicating the minor role of *vll* in Hif regulation in the presence of vhl. However, a striking upregulation of Hif target genes was observed in *vhl-/-;vll-/-* mutants suggesting the redundant role of *vhl* and *vll* in the Hif regulation.

Previously we found that one of Hif hydroxylases, *phd3*, is a direct target of Hif with HREs identified in its promoter [27, 28], and its expression most dramatically reflects the activity of Hif signaling. Accordingly, we used a BAC engineered *phd3* reporter line that expresses EGFP under the *phd3* promoter as a tool to measure the level of HIF activation [26]. As expected, the *phd3* transgenic *vhl* homozygous mutant embryos expressed a very high level of EGFP reflecting Hif activation. The expression of EGFP in *vll-/-; Tg(phd3::EGFP)^i144^* mutant embryos, however, was indistinguishable from that of wild type. When we examined the *vhl-/-;vll*-/-; *Tg(phd3::EGFP)^i144^* double mutants, there was augmented expression of EGFP transgene in comparison to that in *vhl-/-* single mutants (Fig. 1C), confirming the redundant role of *vll* and *vhl* in Hif regulation.

### 2. Zebrafish *vll*-/- mutant fish are more susceptible to loss of heterozygosity (LOH) in the *vhl* locus

Interestingly, in heterozygous *vhl+/-* mutants, the expression of *Tg(phd3::EGFP)^i144^* is very low whereas *vhl-/-* homozygous mutants express a very high level of EGFP, indicating that one copy of Vhl is sufficient to suppress the expression of Hif in the *vhl+/-* heterozygous mutants. While we were examining the *vhl+/-;vll-/-* mutant adult fish, we came across very bright EGFP expressing cells in the mutant fish that were not observed in *vhl+/-;vll+/+* fish. This meant that there was an increased rate of spontaneous LOH of *vhl* locus in the *vll*-/- mutants in comparison to that in wild type fish (Fig. 2A and B).

**Fig. 2.**
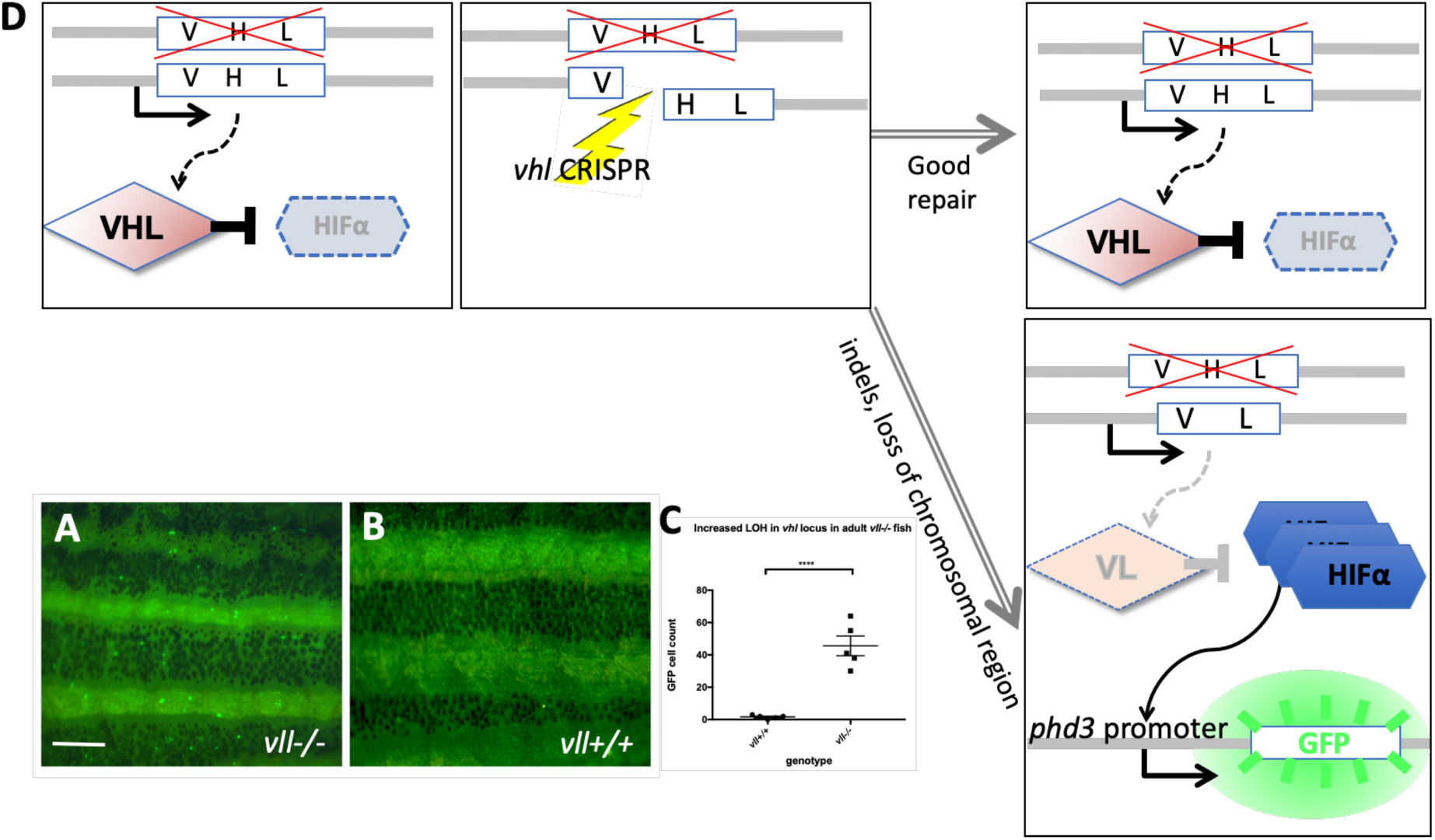
The adult *vll-/-* mutant fish are more susceptible to LOH in the *vhl* locus. A and B. When we examined 6 months old *vll-/-* mutant fish, there was an increased number of spontaneous LOH in the *vhl* locus in *vll-/-* mutant fish in comparison to *vll+/+* wild type fish. The increased LOH in *vll-/-* mutant fish was quantified in C. \*\*\*\**p<0.0001*, unpaired t-test. D. Schematic diagram illustrating the principle of our reporter system that if DNA damage is introduced into the remaining wildtype *vhl* allele, for instance by a CRISPR, in the *vhl+/-:Tg(phd3::EGFP)^i144^* fish in response to the genotoxic stress and the damage is well repaired and no frameshift is introduced, the cells will remain EGFP negative. However, if the DNA damage introduced into the wild type allele is not repaired properly, the cells will lose heterozygosity and express very bright EGFP expression. Scale bar: 1mm

From this observation, we realized that *vhl+/-* heterozygous *Tg(phd3::EGFP)^i144^* transgenic fish could be an excellent tool to study genomic instability. When the cells experience any genotoxic stress which introduces strand breaks or other damage in the DNA of the *vhl* locus, the cells that repair the damage perfectly, e.g. by HR, will remain EGFP negative. On the other hand, if, for instance, a DNA break is repaired by mechanisms that introduce errors or if there are defects in the DNA repair pathways, the cells will lose the remaining wild type allele of *vhl* and will express a very high level of EGFP (Fig. 2C). Therefore, we reasoned that *vll* might be required for maintaining genomic stability and the role of human VHL in DNA repair might be conserved in zebrafish *vll*.

### 3. Zebrafish *vll-/-* mutant embryos increase LOH at *vhl* locus in response to ionizing irradiation

Using *Tg(phd3::EGFP)^i144^* as a tool to study the genomic instability, we treated *vll-/-* mutant embryos with a high level of X-ray to introduce DSBs. We hypothesized that if *vll*, like the human *VHL* gene, is required for the DNA DSB repair by HR upon DNA damage the repair of irradiation induced DNA damage in the *vhl* locus in *vll-/-* mutants would not be as efficient as in wild type embryos, and there will be an increase in the number of EGFP positive *vhl-/-* cells in the *vll-/-* mutant. We irradiated *vhl+/-;vll-/-*; *Tg(phd3::EGFP)^i144^* transgenic embryos with X-rays at 1dpf and examined the LOH at *vhl* locus at 5dpf. Its LOH rate was compared with that of irradiated *vhl+/-;vll+/+* embryos. We found that the LOH rate in the *vhl* locus in the *vll-/-* mutant embryos was much higher than that in *vll+/+* wild type embryos indicating the role of *vll* in accurate DNA repair (Fig. 3A).

**Fig. 3.**
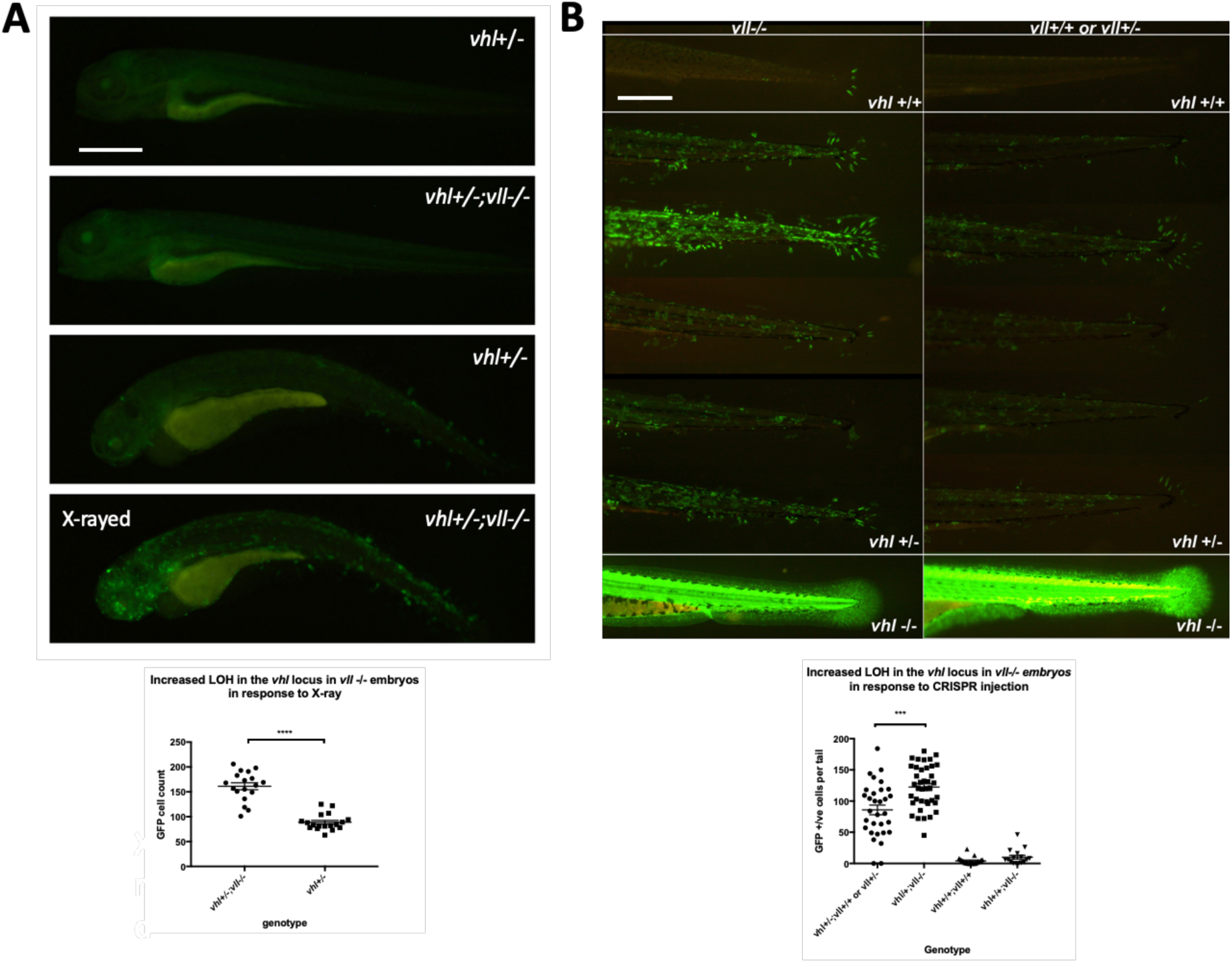
There is an increase in LOH in *vll-/-* embryos in response to X-ray treatments and CRISPR injection. A. There was an increased LOH in the *vhl* locus in *vll-/-* embryos in response to X-ray treatments. \*\*\*\**p<0.0001*, unpaired t-test. B. Injection of gRNA/Cas9 against *vhl* locus induced increased LOH at the *vhl* locus in *vll-/-* embryos in comparison to *vll+/+* wild type embryos. \*\*\**p<0.001*, one way ANOVA. Scale bars: 0.5mm

Next, we used the CRISPR/Cas9 system that specifically targets the *vhl* locus, to introduce a targeted DNA DSBs in *vhl*. Using EGFP expression in the *Tg(phd3::EGFP)^i144^* as a readout for LOH, we injected a single gRNA against *vhl* into 1-cell stage *vhl+/-;vll-/-* embryos and examined the LOH at 5dpf with the equally injected *Tg(phd3::EGFP)^i144^*; *vhl+/-;vll+/+* embryos as their control. Similar to the results from X-ray treatment, there was an increased number of EGFP positive cells in the *vll-/-* mutants in comparison to *vll+/+* wild type embryos, indicating that accurate DNA repair is defective in the *vll* mutant (Fig. 3B).

Loss of heterozygosity can be due to various mechanisms either local mutations or more significant changes like loss of an entire chromosomal region. We investigated efficiency of mutagenesis with CRISPR/Cas9 system that targets not only *vhl* but also another gene, Androgen Receptor (*AR*). We injected guide RNAs targeting *AR /vhl* into *vll*-/- mutants and wild type embryos to quantify mutagenesis rates in the *vll-/-* mutants in comparison to wild type embryos. After the injection, genomic DNAs were extracted and PCR was performed to amplify the sequences around the target sites. We carried out deep sequencing of the PCR products from injected samples. This revealed that there was no significant increase in the mutation frequencies, or in the size of indels in the *vll-/-* mutants in both *AR* and *vhl* targeting CRISPR injected embryos (Supplementary data Fig.1). This suggests that increased LOH in the *vll-/-* embryos might be due the loss of larger regions of the chromosome or complex rearrangements of chromosomes that cannot be detected by PCR methods [29].

To exclude the possibility that *vll* protects embryos from the DNA damage induction rather than it being involved in DNA repair, we examined the expression of *γ*H2AX which is one of the earliest responses to DNA damage after the irradiation. We irradiated *vhl+/-* and *vhl+/-;vll-/-* embryos at 1dpf and fixed them immediately after the X-ray treatment. The fixed embryos were examined for their *γ*H2AX foci formation. This revealed that embryos of all genotypes showed equally distributed formation of *γ*H2AX foci after the X-ray treatment suggesting that DNA damage was equally introduced in all genotypes (Supplementary Fig. 2).

### 4. *vhl-/-;vll*-/- double mutants are protected from X-ray induced apoptosis

We wondered whether *vhl* and *vll* have functional redundancy in their DNA damage repair role, and wanted to investigate the possibility that *vhl* might play a major role in the DNA repair, in the same way as in their Hif regulation. Unfortunately, however, we cannot use our *Tg*(*phd3::EGFP)^i144^* line as a tool to study the role of *vhl* in genomic instability because all the cells in the *vhl* mutants already lack both alleles of wild type *vhl* and express a very high level of EGFP. Therefore, we decided to examine the sensitivity of *vhl-/-;vll-/-* double mutants to genotoxic stress compared with that of *vll-/-* mutant. We reasoned that if *vhl* and *vll* are both required for the DNA repair, the double mutant would be extremely sensitive to the genotoxic stress.

We pair mated *vhl+/-;vll-/-;Tg(phd3::EGFP)^i144^*fish to expose *vhl-/-;vll-/-* double mutants to X-ray treatment. When the collected embryos were irradiated at 1dpf and examined at 5dpf, we found that around a quarter of embryos were extremely well protected from the X-ray induced cell death. We separated these protected embryos from the rest and examined them under the fluorescent microscope. Surprisingly, it turned out that all the protected embryos were *vhl-/-;vll-/-* double mutants with a very high level of EGFP expression (Fig. 4A-D). This was in contrast to our speculation that double mutant would be extremely sensitive to the genotoxic stress. TUNEL staining confirmed a much reduced number of apoptotic cells in the double mutants in comparison to their siblings (Fig. 4E, F).

**Fig. 4.**
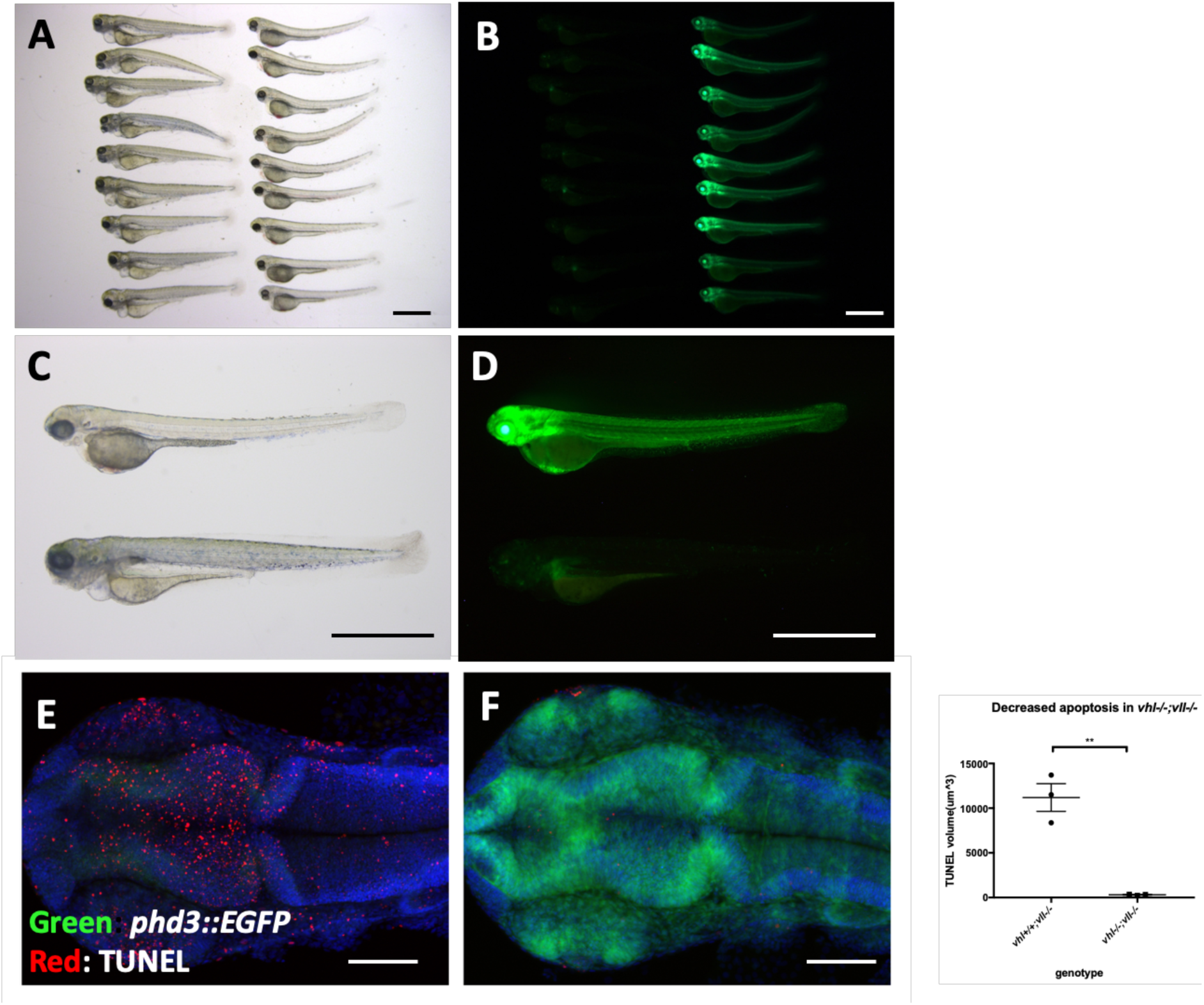
*vhl-/-;vll*-/- double mutants are protected from X-ray induced apoptosis. A-D. The embryos from *vhl+/-;vll-/-* pair mating were collected and the embryos were exposed to X-ray treatment at 1dpf. Then the treated embryos were examined at 5dpf. There were clear differences in response to X-ray treatment; majority of embryos were severely affected by X-ray treatment whereas around 25% of the embryos were well protected without much sign of cell death. We classified these embryos and looked at them under the fluorescent microscope. This revealed that all the protected embryos were EGFP positive *vhl-/-;vll-/-* double mutants. E and F. The embryos were collected from the *vhl+/-;vll-/-* pair mating, and then exposed to X-ray at 1dpf. The embryos were fixed 3 hours post X-ray and we analysed the cell death in the head region with TUNEL. This showed extremely decreased TUNEL positive cells in the *vhl-/-;vll-/-* double mutants in comparison to their siblings. \*\**p<0.01*, unpaired t-test. Scale bars: 1mm (A-D) and 0.2mm (E&F)

### 5. The Hif activation protects embryos from genotoxic stress induced apoptosis

We then wanted to identify what makes *vhl-/-;vll-/-* double mutants resistant to the X-ray treatment. The most obvious effect of the double mutation is upregulation of Hif signaling. Therefore, we sought to upregulate Hif signaling independently of Vhl and Vll to test whether Hif upregulation could protect embryos from cell death induced by genotoxic stress. We employed a cancer therapeutic reagent Camptothecin (CPT), which induces DSBs in replicating cells [30–32]. Embryos from *vhl+/-;vll-/-* pair mating were treated with 20nM CPT at 32hpf overnight. The CPT treated embryos were examined for their sensitivity to CPT at 5 dpf. All the embryos treated with CPT, except *vhl-/-;vll-/-* double mutants that express a high level of EGFP, died by 5dpf with severe cell death (Fig 5A and B) confirming our previous observation. Further, when a parallel batch of embryos were treated with Hif activator (JNJ-42041935) for 8 hours prior to CPT treatment, all the embryos, regardless of their *vhl* genotype, were protected from CPT treatment induced death, suggesting that the protection of *vhl-/-;vll-/-* double mutants from CPT treatment was indeed due to elevated HIF activation (Fig. 5C and D). To confirm these data further, we treated wild type embryos with CPT at 20nM, with or without pre-treatment with Hif activator. This revealed that all the wild type embryos treated with CPT alone died by 5dpf (Fig. 5E), whereas the majority of embryos survived when they were treated with HIF activator prior to CPT treatment (Fig. 5F). Chemical Hif activators function by inhibiting cellular hydroxylases and could therefore have additional effects. To further show that the protection of the embryos is due to upregulated Hif, we injected constitutively active forms of Hif1α and Hif2α to induce a high level of Hif activation, instead of chemically activating Hif. This also showed that all the Hif injected embryos were protected from CPT induced apoptosis, confirming the specificity of chemical Hif activator (Fig 5G, compare to E).

**Fig. 5.**
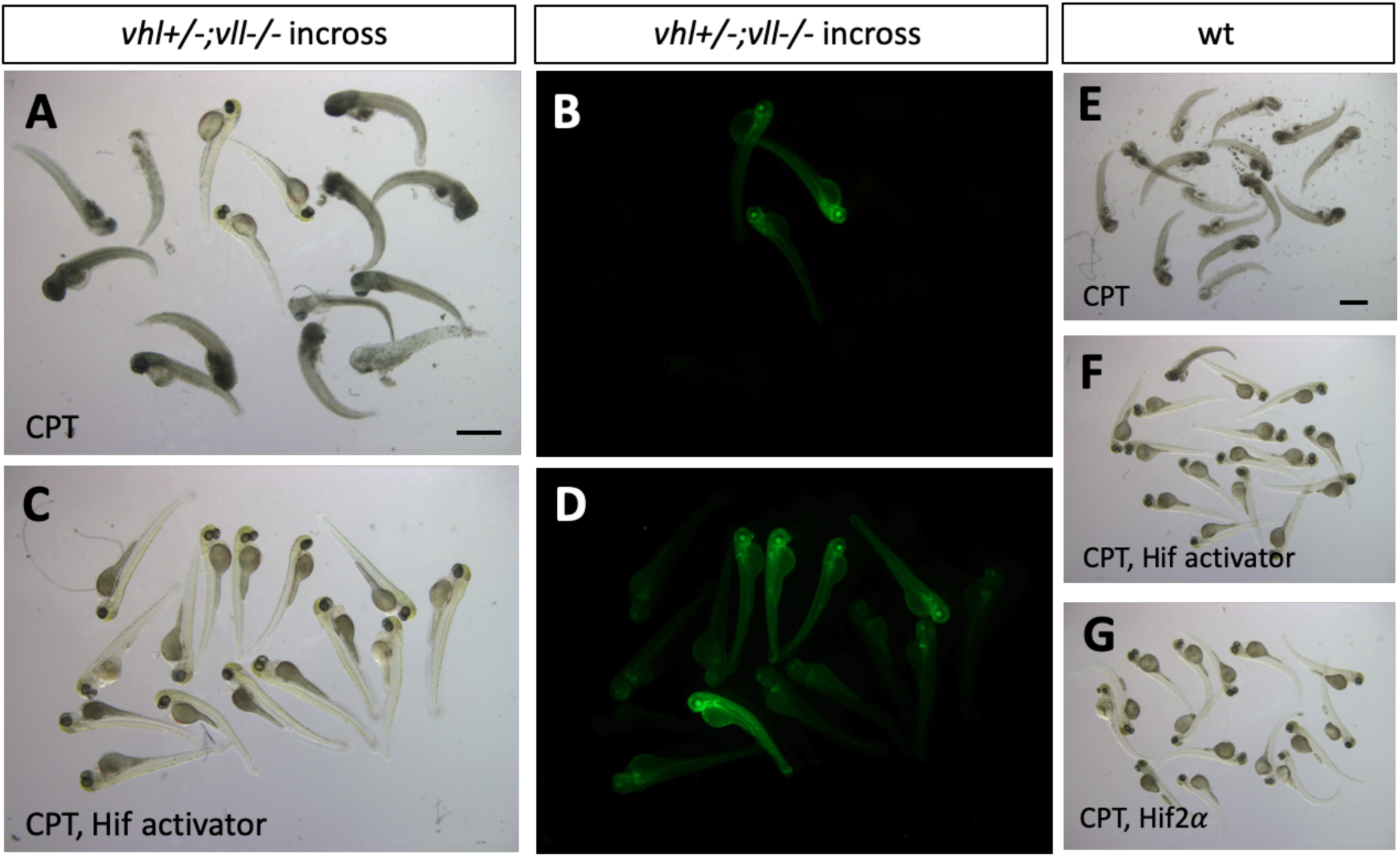
Hif activation protects embryos from CPT induced apoptosis. A and B. The embryos from *vhl+/-;vll-/-* pair mating were treated with 20nM CPT. Majority of embryos did not survive until 5dpf except a few embryos. All the embryos that survived were EGFP positive *vhl-/-;vll-/-* double mutants. C and D. When the embryos were treated with Hif activator for 8 hours prior to the CPT treatment, all the embryos became EGFP positive and survived until 5dpf. E. When wild type embryos were treated with CPT, the embryos did not survive at 5dpf. F. However, most of embryos that are treated with Hif activator prior to CPT treatment survived at 5dpf. G. The embryos injected constitutively active Hif2α also survived at 5dpf. Scale bars: 1mm

### 6. p53 expression is decreased in the *vhl-/-;vll-/-* double mutants and in *vhl-/-* mutants in comparison to their siblings

Previously, a direct role of VHL in p53 stabilisation and transactivation has been reported, with diminished p53 protein levels and activity in the absence of *vhl* [33]. Since we observed reduced apoptosis in the *vhl-/-;vll-/-* mutants and Hif activated embryos in response to genotoxic stress, we questioned whether the reduction in apoptosis in these embryos is due to the downregulation in p53 expression. According to the previous studies in zebrafish and cell culture systems, full length p53 is induced immediately in response to X-ray treatment to remove the cells with excessive DNA damage beyond repair and ensure the cells do not progress through the cell cycle and proliferate. A short form (*Δ*113p53) that is a direct target of full length p53 and transcribed from the C-terminus of intron 4 of full length p53, is induced much later than the full length p53 to promote DNA DSB repair and to antagonise the pro-apoptotic activity of full length p53 [34].

Given to the suppressed apoptosis phenotype observed, we speculated that full length of p53 would be downregulated in the *vhl-/-;vll-/-* double mutants. We *γ*-rayed embryos from *vhl+/-;vll-/-* pair mating and performed q-PCR against full length p53 and the short form of p53. There was a dramatic reduction in both isoforms of p53 expression in the *vhl-/-;vll-/-* mutant embryos in comparison to their siblings. p53 suppression in the *vhl-/-;vll-/-* mutants is likely to occur due to the role of Hif in p53 regulation rather than the direct role of Vhl in p53 regulation as Roe *et al*. suggests, since Hif activator treated embryos also have reduced apoptosis.

To investigate if a reduced p53 level in the *vhl-/-;vll-/-* double mutants may suffice to suppress apoptosis in the mutants, we investigated the effect of p53 downregulation on the apoptosis, using p53 morpholino injection. When we injected p53 morpholino into the embryos that were collected from *vhl+/-;vll-/-* pair mating and treated them with 10nM CPT, we observed all the injected embryos were protected from CPT induced apoptosis regardless of their genotype (Fig. 7B), in contrast to uninjected controls (Fig. 7A) in which only the *vhl-/-;vll-/-* double mutants are protected from CPT treatment. These data are consistent with the idea that double mutants can be protected from genotoxic stress due to a low level of p53 expression. Whether Hif activation leads to suppression of apoptosis by additional mechanisms other than downregulating p53 should be further investigated

**Fig. 6.**
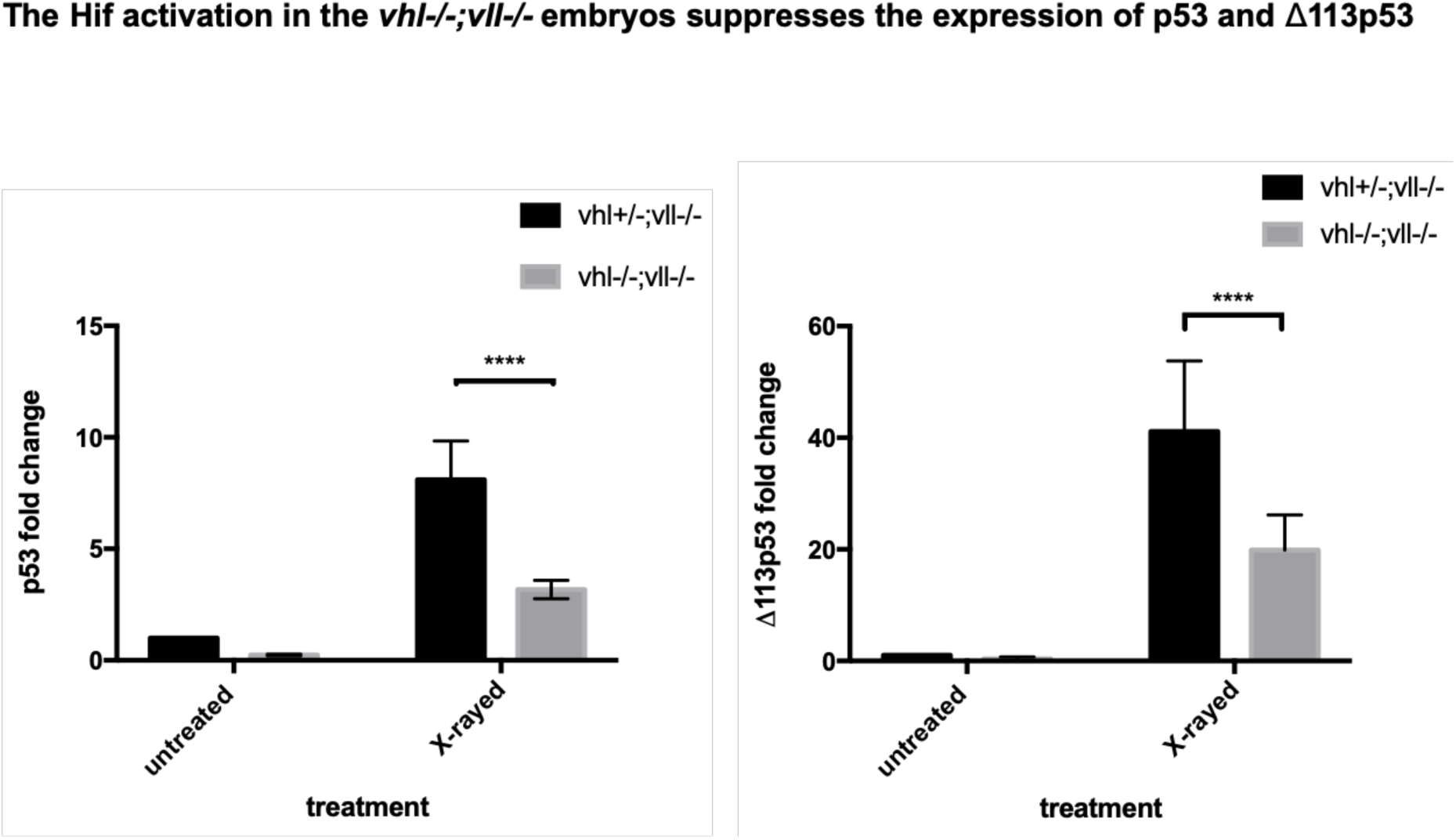
p53 expression in the *vhl-/-;vll-/-* double mutants is downregulated in comparison to that in their siblings. Embryos from *vhl+/-;vll-/-* pair mating were collected and irradiated at 2 dpf with *γ*-ray at 20 gray. q-PCR was performed 24 hours post *γ*-ray to quantify the full length of p53 and short form of *Δ*113p53. This revealed that there was a significant decrease in both forms of p53 expression in the *vhl-/-;vll-/-* embryos in comparison to their siblings. \*\*\*\**p<0.0001*, two way ANOVA test. The statistical analysis was performed with ddCt values.

**Fig. 7.**
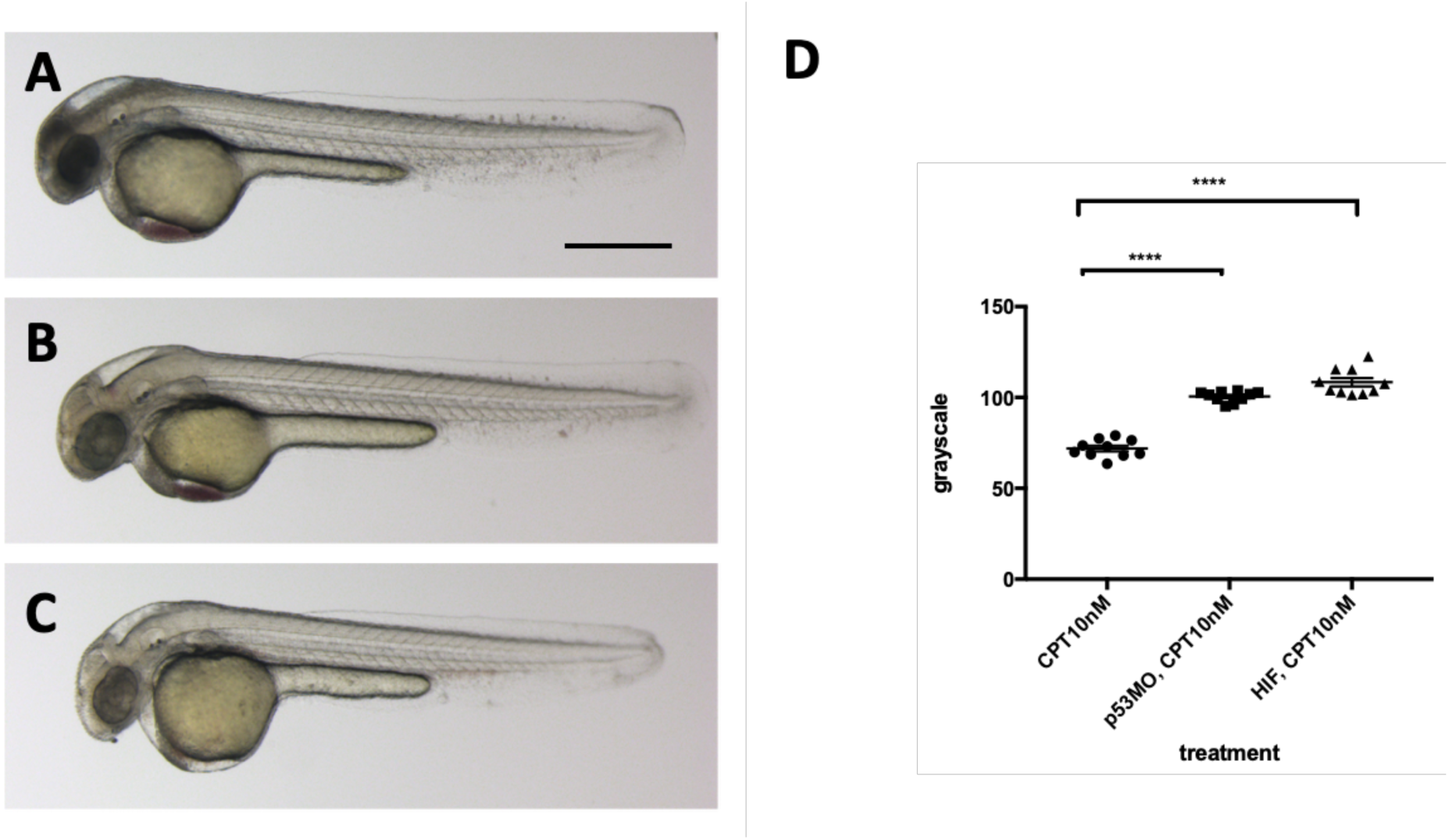
p53 knock down by p53 morpholino injection protects embryos from CPT induced apoptosis. A. Embryos collected from *vhl+/-;vll-/-* pair mating were treated with 10nM CPT. This resulted in severe cell death in the CPT treated embryos with the exception of GFP positive *vhl-/-;vll-/-* double mutants. B. On the contrary, when p53 morpholino was injected into the embryos prior to 10nM CPT treatment, all the embryos were protected from CPT induced apoptosis. C. The similar protection was also observed in the embryos treated with Hif activator prior to CPT treatment. Only GFP negative embryos were imaged and quantified for this experiment. \*\*\*\**p<0.0001*, one-way ANOVA. Scale bar: 0.5mm

### 7. The Hif activation protects embryos from genotoxic stress induced DNA damage

The results clearly demonstrate the role of Hif in the suppression of apoptosis, probably through downregulation of p53. There are two scenarios that can explain these observations, either lack of p53 activation prevent cells with severe DNA damage from undergoing apoptosis, and consequently these cells with accumulated DNA damage will contribute to the cell population of later embryos, or alternatively HIF might have a truly genoprotective role, improving the resilience of the genome to DNA damage, thereby reducing the p53 activation. To test whether HIF activation plays a role in the genoprotection, we decided to test its effect on the increased DNA damage in the *vll* mutants. We treated *vll-/-* mutants with HIF activator for at least 2 hours prior to X-ray treatment at 24hpf. These embryos were heterozygous for *vhl* and carried *Tg(phd3::EGFP)^i144^* to allow the detection of LOH levels for *vhl,* as described previously. We would expect that if cells with damaged DNA fail to undergo apoptosis due to lack of p53 function, more cells with LOH might be visible per embryo, but if Hif has a role in protecting cells from DNA damage, the LOH rate will be reduced. The embryos were examined for their LOH at 5dpf. This revealed that *vll-/-* mutant fish treated with Hif activator had significantly decreased LOH (Fig. 8 B and D; quantified in E), suggesting that Hif itself has a role in the protection of embryos from DNA damage and that the role of Vll itself in DNA repair is likely to be Hif independent. In addition, Hif activator treated embryos looked a lot healthier than untreated embryos, which is consistent with the results from CPT treatment. To quantify the latter we used embryonic eye size as an easily measurable “surrogate” for overall health, and found it to be significantly increased (compare Fig. 8A and C; quantified in F).

**Fig. 8.**
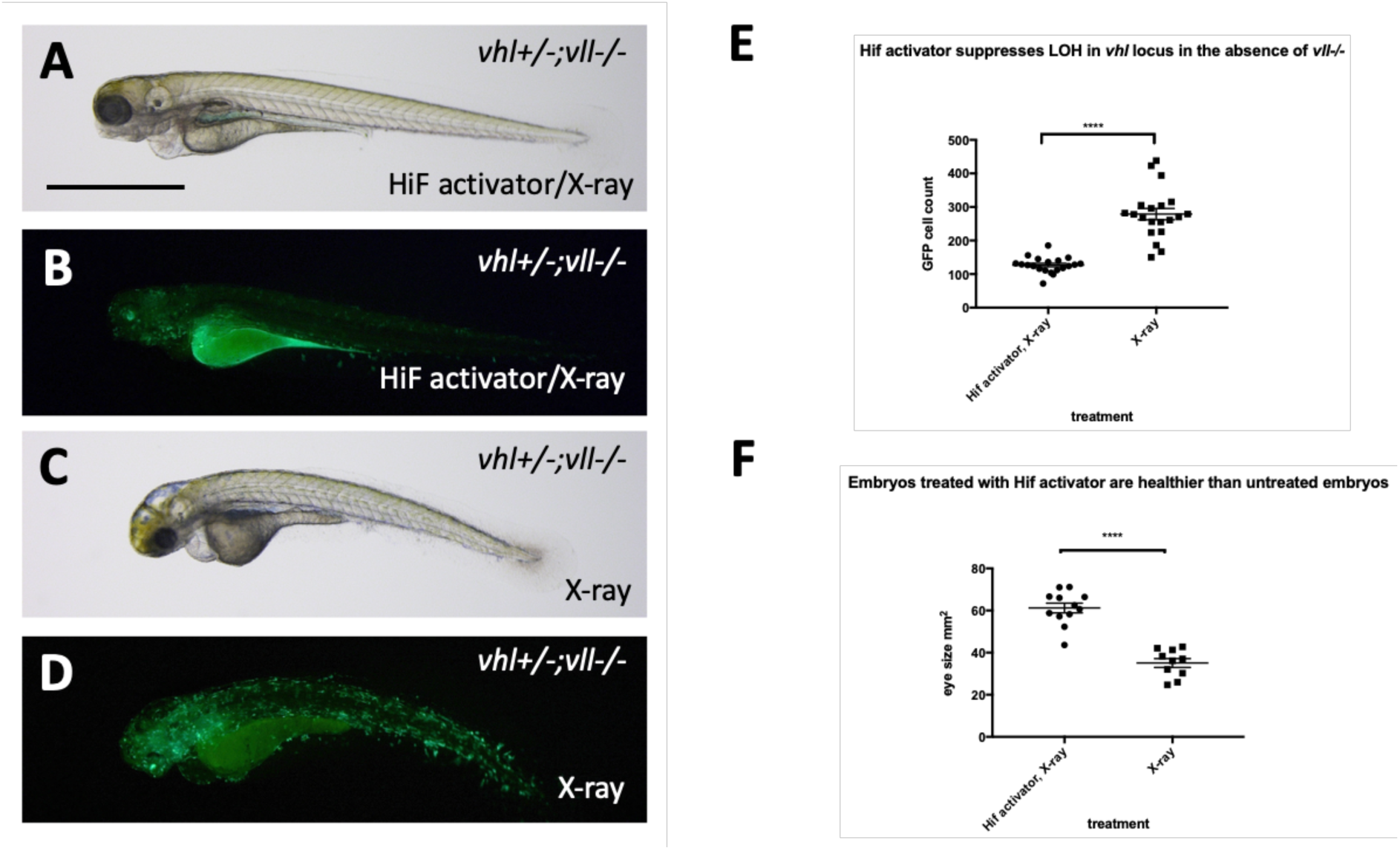
Hif activator treatment protects embryos from X-ray induced DNA damage. A-D We investigated the effect of Hif activation on the increased LOH in *vhl* locus in *vll-/-* mutant embryos. When embryos were treated with Hif activator prior to X-ray treatment, the increased LOH in the *vhl* locus in *vll-/-* embryos was significantly reduced (B) in comparison to *vll-/-* embryos that were exposed to X-ray without Hif activator treatment (D). E. The effect of Hif activator on the LOH suppression in the *vhl* locus was quantified. \*\*\*\**p<0.0001*, unpaired t-test. Hif activator treated *vll-/-* embryos looked more healthier (A) than untreated embryos (C). F. The size of eyes was measured and quantified in the Hif activator treated and untreated embryos. \*\*\*\**p<0.0001*, unpaired t-test. Scale bar: 1mm

Then we asked whether downregulation of p53 has any effect on the DNA repair function of Hif. We injected p53 morpholino into the embryos collected from *vhl+/-;vll-/-;Tg(phd3::EGFP)^i144^* pair mating and treated them with CPT to examine the effect of p53 downregulation on the LOH formation in *vhl* locus. We counted the number of EGFP positive cells in the *vhl+/-;vll-/-* embryos that are induced by CPT treatment and compared with the number of EGFP positive cells in the p53 morpholino injected *vhl+/-;vll-/-* embryos. There was no significant difference in the number of EGFP positive cells between p53 morpholino injected and uninjected *vll-/-* embryos, indicating the role of HIF in the promoting DNA repair is through p53 independent mechanism (Fig. 9 A and B).

**Fig. 9.**
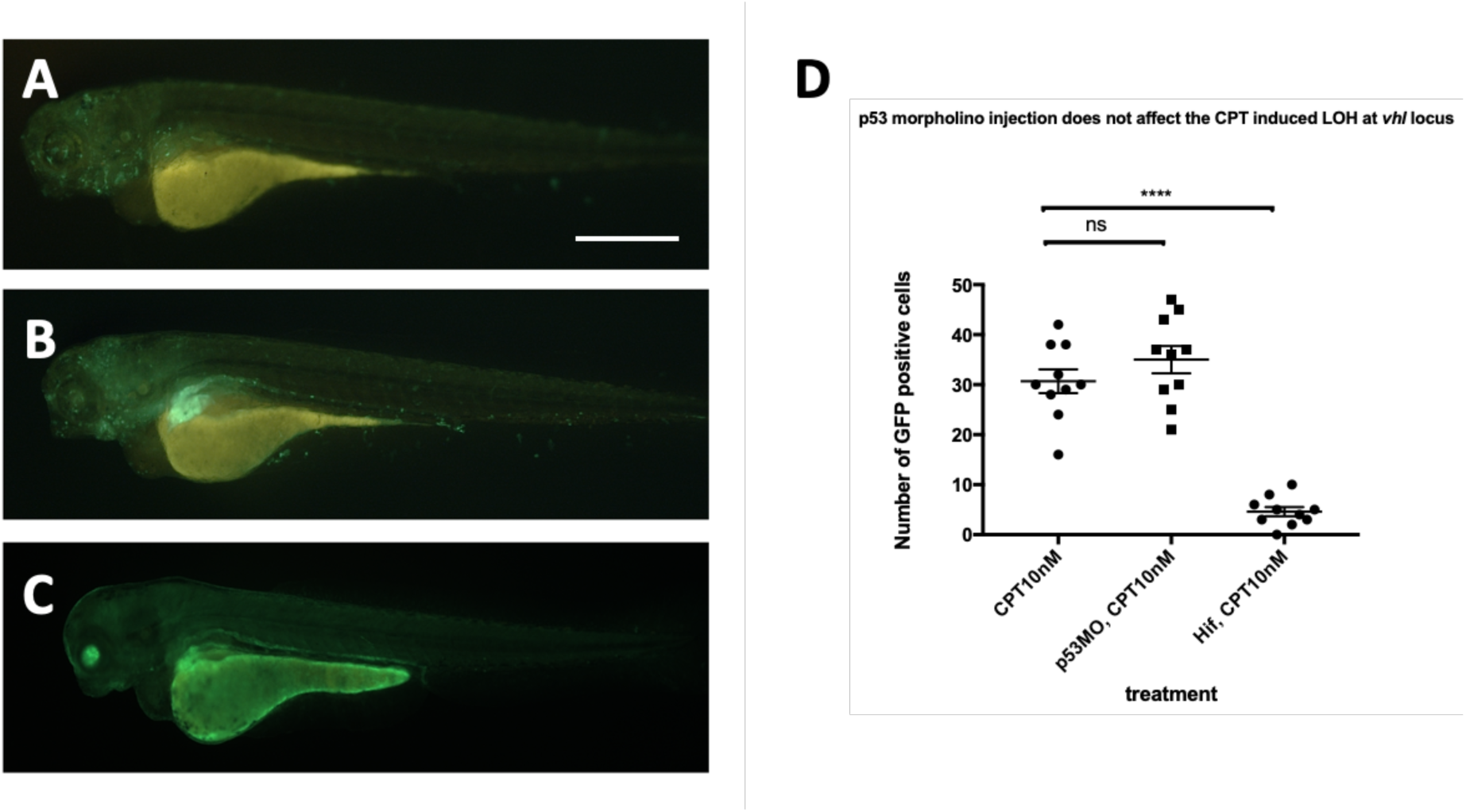
p53 knock down by p53 morpholino does not affect the number of LOH at the *vhl* locus induced by CPT. A. The embryos collected from *vhl+/-;vll-/-* pair mating were treated with 10nM CPT. There was GFP positive cells throughout the CPT treated embryos indicating the LOH in these cells. B. A similar number of GFP positive cells were observed in the p53 morpholino injected embryos suggesting that the CPT induced LOH formation in these embryos is independent of p53 level. C. On the contrary, the treatment of Hif activator prior to CPT treatment abolished the LOH formation in the embryos, signifying the genoprotective role of Hif against CPT treatment. ^ns^ *p>0.05*, \*\*\*\**p<0.0001*, one-way ANOVA. Scale bar: 0.5mm

### 8. *brca2-/-* mutants are sensitive to X-ray treatment induced cell death and an elevated Hif level in the *vhl-/-* mutants alleviates the cell death induced by mutation in *brca2*

As the precise relationship of Vll to DNA repair is still very poorly understood, we sought to look at the effect of Hif on well-established DNA repair gene mutations. Firstly, we created a *brca2* mutant allele using the CRISPR/Cas9 system. The *brca2* mutant allele for this experiment contained an 83bp insertion at the amino acid 445 (exon 10) position introducing premature stop. When embryos from *brca2+/-* pair-mating were treated with X-rays, we found that there was increased cell death in homozygous *brca2-/-* embryos, especially in the central nervous system (CNS) as previously reported [35]. In contrast, the least affected embryos were all either *brca2+/-* or *brca2+/+* (Fig. 10A). We confirmed the increased apoptosis in the *brca2-/-* embryos in comparison to their siblings by Acridine Orange staining (Fig. 10B).

**Fig. 10.**
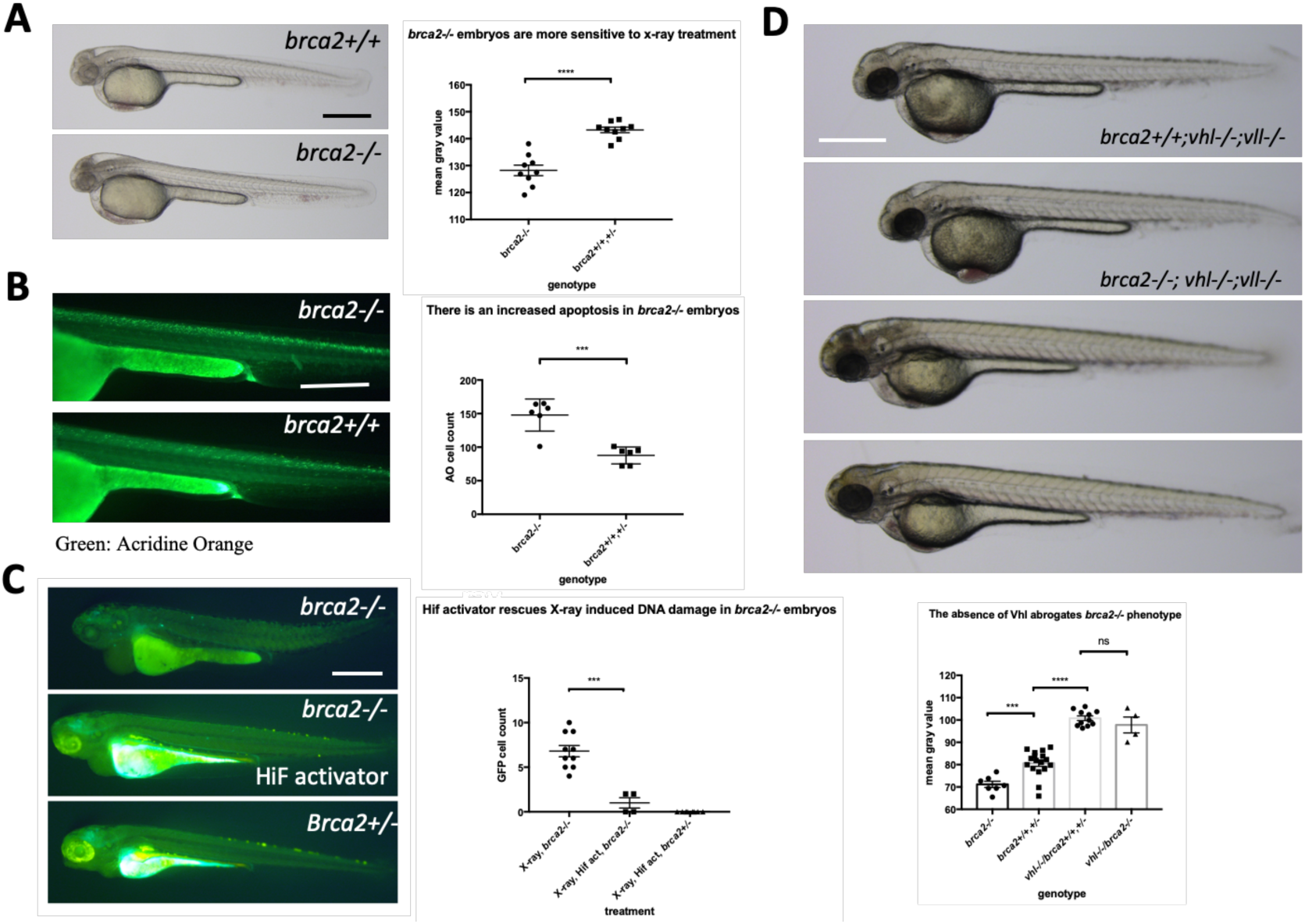
The elevated Hif level in the *vhl-/-* mutants alleviates cell deaths induced by the mutation in *brca2*. A. *brca2-/-* embryos were more sensitive to X-ray treatment in comparison to their siblings demonstrating the increased cell death in CNS. \*\*\*\**p<0.0001*, unpaired t-test. B. The Acridine Orange staining confirmed the increased apoptosis in the *brca2-/-* embryos. \*\*\**p<0.001*, unpaired t-test. C. X-ray treatment induced a few EGFP positive cells in the *brca2-/-* embryos, indicating the loss of both *vhl* alleles in these embryos. On the other hand the EGFP positive cells were hardly observed in their siblings. The Hif activator treatment suppressed the appearance of EGFP positive *vhl* mutant cells in the *brca2-/-* embryos. \*\*\**p<0.001*, one way AVOVA. D. *brca2-/-* embryos were more sensitive to X-ray treatment in comparison to their siblings. However, when the *vhl* -/-;*vll*-/- mutations are introduced in the *brca2-/-* embryos, increased apoptosis in the CNS of *brca2-/-* embryos was suppressed and the *brca2-/-* embryos were indistinguishable from their siblings. \*\*\**p<0.001*, \*\*\*\**p<0.0001*, ^ns^ *p>0.05*, one way ANOVA. Scale bars: 0.5mm

We then proceeded to check whether elevated Hif activation seen in the *vhl-/-;vll-/-* mutants can also suppress the increased sensitivity of *brca2-/-* embryos to genotoxic stress. We generated *brca2+/-;vhl+/-;vll-/-* mutants. The embryos from *brca2+/-;vhl+/-;vll-/-* in cross were treated with X-rays at 1dpf and examined for the cell death in the CNS 2 days post X-ray. This revealed that the increased apoptosis in the *brca2-/-* mutant was suppressed by the presence of *vhl-/-;vll-/-* and that *brca2-/-* embryos were indistinguishable from *vhl-/-;vll-/-* embryos, suggesting that the elevated Hif can strongly suppress the sensitivity of *brca2-/-* embryos to X-ray induced apoptosis (Fig. 10D).

In addition, we examined the effect of the loss of *brca2* in the rate of loss of *vhl* alleles in the *brca2-/-;Tg(phd::3EGFP)^i144^*embryos by X-ray treatment. When we exposed the embryos from *brca2+/-;Tg(phd::3EGFP)^i144^* pair mating to a high dose of X-ray, there were a few cells expressing a high level of EGFP in *brca2-/-* embryos, indicating that these cells had lost both *vhl* wild type alleles, in contrast to their siblings in which there were hardly any EGFP positive cells. When these embryos were pre-incubated with Hif activator before the X-ray treatment, the number of EGFP positive cells in the *brca2-/-* embryos was profoundly reduced, proving the role of Hif in the protection of *brca2 -/-* embryos from the X-ray induced DNA damage (Fig. 10C).

### 9. Hif activation abrogates the sensitivity to CPT induced cell death in ATM inhibitor treated embryos

Another fundamental regulator of the DNA damage response is ATM (ATM serine/threonine kinase), and ATM deficient cells are highly sensitive to genotoxic stresses [36]. Patients with mutations in ATM develop A-T syndrome that are characterised by progressive neurodegeneration, increased risk of cancer development, radiosensitivity and immune system impairment. We wondered whether Hif activator can mitigate the effect of the loss of ATM through its role in the anti-apoptosis and DNA repair. We treated wild type embryos with 10nM CPT with or without pre-incubation with 10μM ATM inhibitor (ATMi). As expected, ATMi treated embryos were highly sensitised to CPT treatment with increased apoptosis in the brain in comparison to ATMi untreated embryos. However, when the embryos were treated with Hif activator in combination with ATMi before CPT treatment, Hif activator treated embryos were well protected from CPT induced cell death (Fig.11 A-C). We also observed the protection of ATMi treated embryos from X-ray induced LOH by Hif activator treatment. ATMi treated embryos were highly sensitised to X-ray treatment with increased LOH at the *vhl* locus in comparison to ATMi untreated embryos. On the contrary, when the embryos were treated with both Hif activator and ATMi, the increased LOH at the *vhl* locus was dramatically reduced (Fig. 11 D-F).

**Fig. 11.**
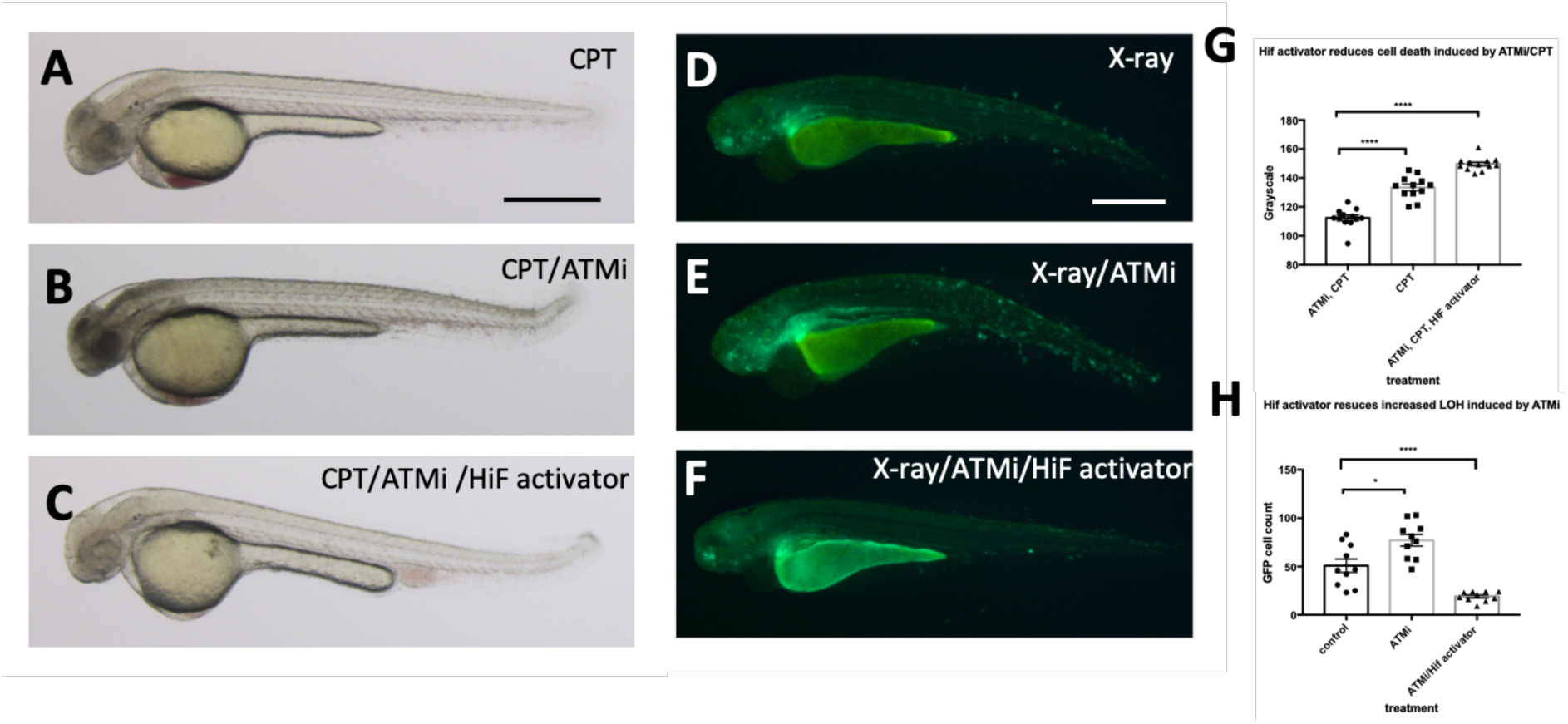
Hif activation abolishes the sensitivity to CPT induced cell death in ATM inhibitor treated embryos. A. The wild type embryos were treated with 10nM CPT to induce cell death in the CNS. B. When the embryos were treated with ATMi before CPT treatment, the embryos were highly sensitised to CPT treatment and induced severe cell death in the CNS. C. When the embryos were treated with Hif activator in combination with ATMi prior to CPT treatment, the embryos were well protected from CPT/ATMi induced cell death in CNS. The quantification of the data is shown in G. \*\*\*\**p<0.0001*, one way ANOVA. D. The *vhl+/-;vll-/-;Tg(phd3::EGFP*)*^i144^*embryos were treated with X-ray to induce LOH at the *vhl* locus. E. When the *vhl+/-;vll-/-;Tg(phd3::EGFP*)*^i144^* embryos were treated with ATMi before the X-ray treatment, there was a dramatic increase in the LOH in the *vhl* locus in the ATMi treated embryos. F. When the embryos were treated with Hif activator in combination with ATMi before the X-ray treatment, there was a remarkable reduction in the LOH in the *vhl* locus. The quantification of the data is shown in H. \**p<0.05*, \*\*\*\**p<0.0001*, one way ANOVA. Scale bars: 0.5mm

### 10. The HIF activator does not protect human MRC5 cells from X-ray treatment

We were very surprised by the strength of the protective effect that Hif conferred in zebrafish embryos and we were eager to see if it is conserved in cultured human cells. As cancer cells may have defects in p53 or other relevant pathways, we initially chose MRC5 that is the most common human diploid fibroblasts cell line derived from normal lung tissue of a 14 weeks old male fetus. We took MRC5 cells with or without HIF activator (FG4592) at 15uM concentration and the cells were incubated for 3 hours at 37°C followed by X-ray treatment at 5 gray. The cells were fixed 4 hours and 24 hours post irradiation (hpi) and stained with *γ*H2AX to examine the DNA damage foci. Surprisingly, we observed that there was no statistical difference in *γ*H2AX formation between the HIF activator treated group and non-treated group at both time points (Fig.12 A). In our second experiment, we decided to lower the X-ray dose to 2 gray since we could not observe the resolution of *γ*H2AX at 24 hpi from the first experiment and tested an alternative cell line, human osteosarcoma U2OS. U2OS cells were treated with HIF activator (JNJ-42041935) at 100uM concentration and incubated cells for 3 hour before the X-ray treatment at 2 gray. The cells were fixed 30 minutes post irradiation (mpi) and 24 hpi to examine *γ*H2AX formation. When we counted the number of *γ*H2AX foci, we still could not observe the genoprotective effect of HIF activator treatment at both time points (Fig. 12 B).

**Fig. 12.**
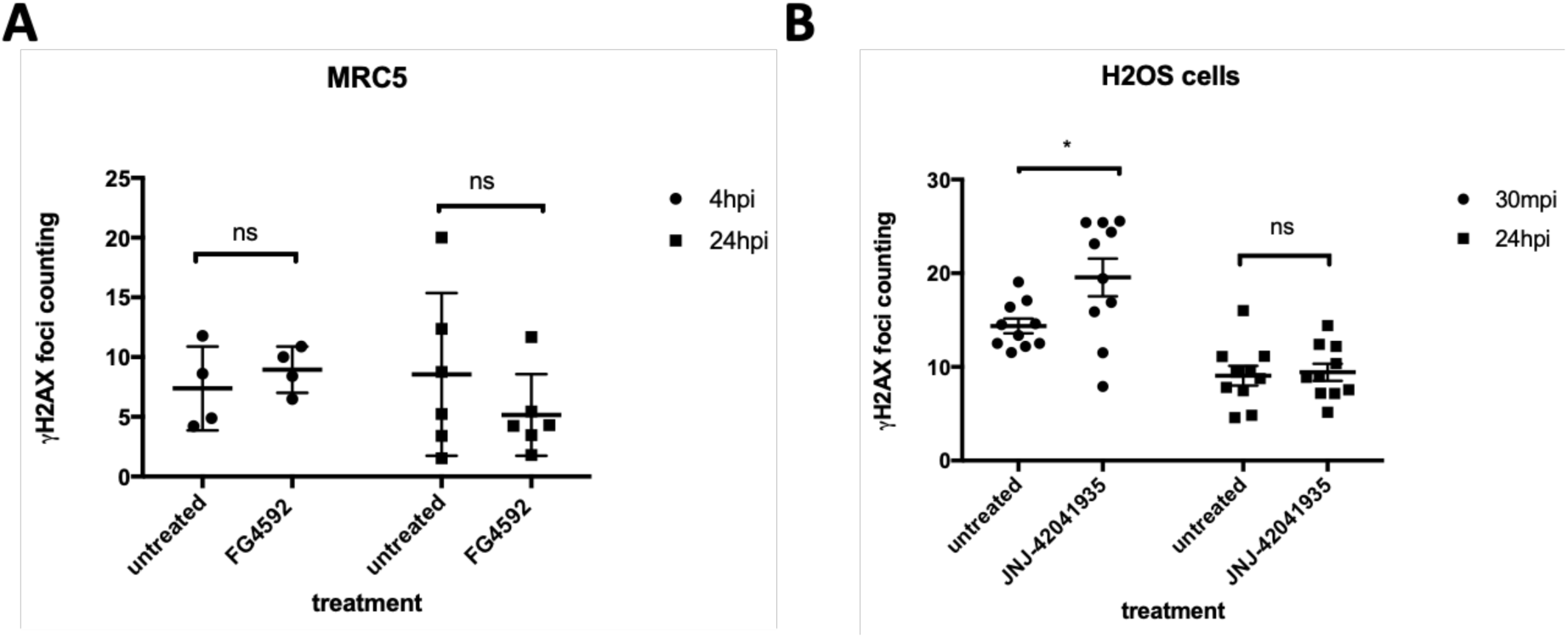
HIF activation does not protect mammalian cells from X-ray induced DNA damage. A. We treated MRC5 cells with HIF activator FG 4592 for 3 hours prior to X-ray treatment at 5 gray. The cells were fixed at 4 hpi and 24 hpi with 4% PFA and stained with *γ*H2AX. This revealed that there was no significant difference in the *γ*H2AX formation between HIF activator treated cells and non-treated cells. ^ns^ *p>0.05*, one way ANOVA. B. The cells were treated with HIF activator JNJ-42041935 for 3 hours prior to X-ray treatment at 2 gray. The cells were fixed at 30 mpi and 24 hpi and stained with *γ*H2AX. At both time points, we could not observe the reduction in *γ*H2AX formation in the HIF activator treated group in comparison to untreated group. At 30mpi, there was even an increase in the number of *γ*H2AX foci in the HIF activator treated group in opposition to our expectation. ^ns^ *p>0.05*, \**p<0.05*, one way ANOVA.

### 11. The Hif activator treatment does not promote colony formation in the MRC5 cells

As an alternative readout for Hif mediated protection to DNA damaging treatment in cell culture, we performed a clonogenic survival assay. The MRC5 cells (4000 cells) were seeded a day before the drug treatment. The cells were then treated with HIF activators with or without ATM inhibitor. 3 hours later, CPT was added either at 10nM or 25nM concentration. The drugs were washed off after 1 hour incubation. The cells were then left to form colonies in the 37°C CO_2_ incubator. Seven days later, the colony formation was examined. There was decreased number of colonies in the CPT/ATM inhibitor treated cells in comparison to untreated control cells. Unlike in zebrafish, however, in the cell culture studies, we could not find any increase in the survival rates in the Hif activator treated cells in comparison to the non-treated cells (Fig.13).

**Fig. 13.**
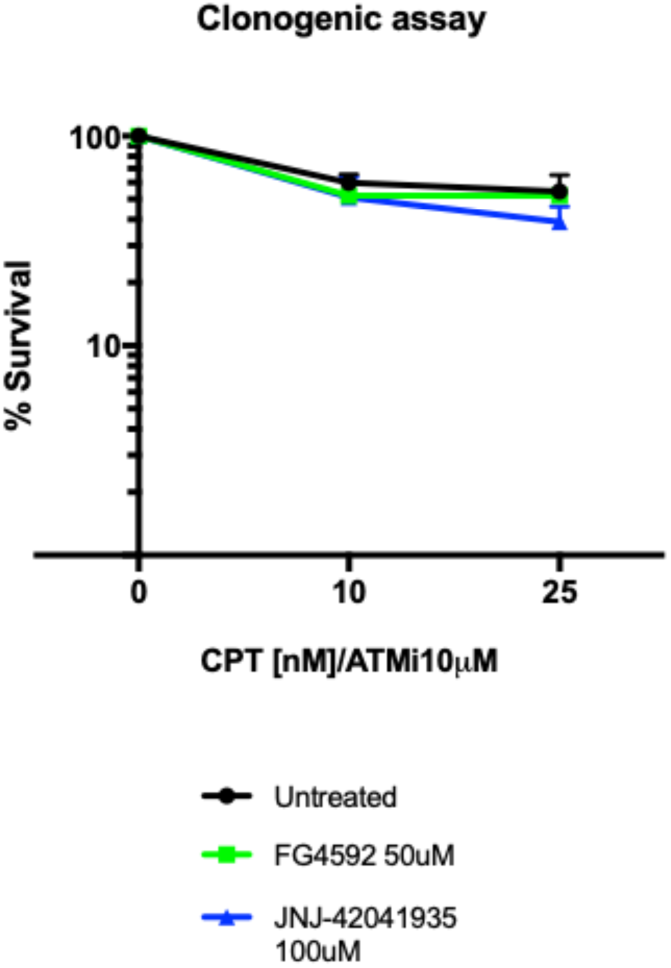
The Hif activator treatment does not affect the survival of MRC5 cells. 4000 MRC5 cells were seeded in 6 well plates and left to grow overnight. The next day, the cells were treated with HIF activator (FG4592 or JNJ-42041935) and ATMi for 3 hours. CPT was added at either 10nM or 25nM concentration for 1 hour and the drugs were washed off. The colony formation was examined 7 days after the drug treatment. CPT/ATMi treatment decreased the survival rates but we could not find any effect of HIF activator on the survival of MRC5 cells. ^ns^ *p>0.05*, unpaired t-test.

## Discussion

The ccRCC is one of the most difficult diseases to treat because of its resistance to chemotherapies and radiotherapies. Although a few targeted therapies have improved the survival rates of ccRCC patients to a certain extent, advanced ccRCC still remains incurable. In addition, unlike some other tumor suppressor proteins, pVHL is a highly multifunctional protein that makes it difficult to design target therapies against. Evidently upregulated HIF in ccRCC provides survival advantages to cancer cells. The activation of HIF, however, although necessary for the ccRCC progression, is not sufficient to initiate tumor development [37–39].

According to the mutator phenotype theory in cancer, the rate of random mutations cannot explain the frequency of mutations in cancer, and the mutations in the genes that are essential for the maintenance of genomic stability, including genes that are involved in DNA repair, have been speculated to increase the mutation rates and drive cancer development [40, 41]. In line with this theory, Metcalfe *et al*. demonstrated that human pVHL is required for the DSBR in the ccRCC cell lines, leading us to speculate that the loss of DNA repair function of VHL might account for the initiation of ccRCC in the VHL patients [7]. We set out to corroborate these data using a whole organismal vertebrate model, zebrafish, to overcome the limitation of the already transformed cancer cell lines for the functional studies for VHL.

We found that in zebrafish, among two VHL homologous genes, the role of VHL in HIF regulation is mainly performed by Vhl, whereas the role of Vll in Hif regulation is very modest. We found that, consistent with the studies in the ccRCC cell lines, *vll* mutant embryos were more susceptible to genotoxic stress and induced increased LOH in our assay with *Tg(phd3::GFP)^i144^* reporter line, indicating the role of VHL in DNA repair is conserved in Vll in zebrafish. However, when we performed deep sequencing around the target sites of CRISPRs against AR and Vhl, the rate of mutation frequencies generated in *vll* mutants was not significantly different from *vll* wild type embryos. The precise reasons for this remain unclear, perhaps the defects that are observable by the LOH assay may consist of more significant changes to the *vhl* containing chromosome, e.g. the loss of a chromosome arm, as is thought to occur in VHL patients [29]. As our next gene sequencing assay is based on PCR amplification, such deletion events go undetected.

We hypothesized that Vhl might have a more dominant role in the DNA repair than its homologous gene Vll, considering its higher homology with human VHL. Unfortunately, we could not use *Tg(phd3::GFP)^i144^*LOH reporter assay to study the LOH in the *vhl* mutants, since all the cells in the *vhl* mutants already express a high level of GFP and there is no wild type *vhl* allele remaining that can be used as “reporter”. We reasoned, however, that *vhl-/-;vll-/-* double mutants will have an increased sensitivity to genotoxic stress, if not only *vll* but also *vhl* is involved in the DNA repair pathway. But in marked contrast to our expectation, we found that *vhl-/-;vll-/-* double mutant embryos were highly protected from the genotoxic stress with much reduced apoptosis, when the embryos were treated with X-ray and CPT.

Considering the prominent upregulation of Hif in the *vhl-/-;vll-/-* double mutants, we then tested whether the elevated Hif in double mutants is responsible for the protection of embryos from the genotoxic stress induced cell death. When we treated embryos with pharmacological Hif activator prior to X-ray and CPT treatment, Hif activation resulted in the protection of embryos from apoptosis induced by the genotoxic stress, even in the presence of intact *vhl* and *vll*. Injection of dominantly active form of Hif into the newly fertilized embryos also resulted in the protection of embryos from the CPT treatment in the same way as pharmacological activation of Hif results, confirming the role of Hif in the providing embryos with protection against genotoxic stress.

In addition, when pharmacologically activated, Hif suppressed the increased LOH in the *vll* mutants. We speculate that this may happen by activating as yet unidentified DNA repair pathways. As a result, in the *vhl-/-;vll-/-* double mutants in which there is a maximal level of increased Hif signaling, the DNA repair defect due to the lack of *vll* was suppressed by the elevated Hif activation and the embryos were well protected from DNA damage and apoptosis. On the contrary, the moderate upregulation of Hif in the *vll* single mutants was not sufficient to suppress the DNA repair defect caused by the lack of Vll function in DNA repair, manifesting the increased LOH in the *vhl* locus in response to ionizing irradiation.

Our data suggest that zebrafish VHL homologues play two opposing roles. On the one hand, they suppress DNA repair by suppressing Hif; on the other, *vll* promotes DNA repair. Therefore, we speculate that the balance of these two opposing roles of VHL might be associated with cancer initiation. HIF activation might initially be able to “mask” effects of the loss of VHL on genomic stability, however when the malignant cells finally form, they will be protected by the strong antiapoptotic function of HIF. Our data is consistent with resistance to the chemo and radio therapies seen in later stage of ccRCC in human through HIF’s role in DNA repair and suppression of apoptosis. We propose zebrafish *vhl-/-;vll-/-* mutants as an excellent platform for the screening for drugs that can interfere with the protection that is conferred by Hif, and therefore overcome resistance.

We discovered that the level of p53 was downregulated in *vhl-/-;vll-/-* double mutants in response to the ionizing irradiation in comparison to their siblings. In addition, when we injected embryos with p53 morpholino, the injected embryos were all protected from CPT induced apoptosis even in the presence of intact *vhl*, suggesting that the decreased level of p53 is responsible for the protection of embryos from apoptosis in the *vhl* mutants, at least partially. This is consistent with *in vitro* studies in which the role of pVHL in the p53 activation has been demonstrated [33]. However, we propose that the downregulated p53 level in the zebrafish *vhl* mutants is due to the elevated Hif, rather than by the Hif independent role of Vhl in regulation of p53. Importantly we found that the downregulation of p53 did not affect the increased LOH in the *vll* mutants, signifying the fact that the role of Hif is not simply in regulating the level of p53, but Hif has a genuine role in the DNA repair. The downregulation of p53 could be the consequence of reduction in DNA damage due to the function of Hif in DNA repair.

Most interestingly, Hif activation also suppressed the increased apoptosis and DNA damage induced by X-ray and CPT treatment in the *brca2* mutant and in the embryos with deficient ATM function. Both BRCA2 and ATM play important roles in the DNA repair pathways in human and their mutations increase the risk of cancer development. In addition, one of the distinctive features of A-T syndrome caused by the deficient ATM function is the defective movement and coordination due to neurodegeneration in cerebellum [42]. Similarly, the mutations in genes that encode proteins in the DNA repair machineries often lead to congenital neurodegenerative disorders in human, highlighting the importance of maintaining the genomic stability in the nervous system. For example, mutations in MRE11, TDP1 and aprataxin (APTX) result in neurodegenerative disorders such as A-T like disease (ATLD) [43], spinocerebellar ataxia with axonal neuropathy (SCAN1) [44], and ataxia-oculomotor apraxia-1 [45, 46] respectively. The efficacy of Hif activators for the suppression of DNA damage and apoptosis in *brca2* mutants and ATM inhibitor treated embryos in our study suggests that HIF activators might provide a possible intervention to slow the progression of both congenital and age related neurodegenerative disorders.

Unfortunately, we could not reproduce the data we generated in the zebrafish in MRC5 and U2OS cells; HIF activation did not protect cells from X-ray induced DNA damage in these cell lines. HIF activator treatment did not affect the survival rates of cells after CPT treatment in the MRC5 cells either. We cannot currently explain why this is the case. We wanted to use the cell lines to gain mechanistic insight with regard to the particular DNA lesions and specific DDR proteins that are affected by HIF activation, taking advantage of abundant resources of antibodies which are available in the cell culture system. Nevertheless, our *in vivo* data are very clear, and it is perhaps an illustration of the importance of using *in vivo* systems to complement cell culture work. Therefore, the limitation of our study is the lack of mechanistic understanding; we are left to raise questions, what specific DNA lesions are protected from Hif activation, what are the downstream target genes of Hif in these processes and how Hif interacts with other proteins and are there any other targets of Hif than p53 that provide survival advantages to the cells lacking the functional Vhl? Given time, we could generate mutants that harbor mutations in the DNA repair pathway, and study how Hif activation can modulate the response of the mutant embryos to the genotoxic stress.

In summary, we discovered that HIF regulation and DNA repair role of human VHL are conserved in zebrafish *vhl* and *vll*. The activated Hif in the *vhl;vll* mutants strongly suppresses DNA damage and apoptosis induced by genotoxic stress. We speculate this could parallel resistance to chemo- and radio therapies in ccRCC, and propose zebrafish *vhl-/-;vll-/-* double mutants as a powerful model for the development of therapeutic reagents to overcome the resistance of ccRCC to chemo- and radio therapies. We found Hif activation suppresses the DNA damage in the *brca2* and *vll* mutant and ATM deficient embryos and also prevent apoptosis. Currently HIF activator is in the phase III clinical trials for the treatment of anemia [47, 48]. Our results will enforce the idea that HIF activation could have a clinical benefit in an increasing number of neurodegenerative disorders that have linked with DNA damage.

## Materials and Methods

### Zebrafish husbandry

All the zebrafish strains used in this paper are maintained on 10/14 hours dark/light cycle at 28°C according to the UK home office regulation at the University of Sheffield.

### Mutagenesis

We generated *vll* mutants by zinc finger nucleases (ZFNs) purchased from Sigma-Sangamo Biosciences. A pair of ZFNs designed to bind the following sequences (in capitals) showed good activity when RNA encoding these was injected into embryos, and the efficiency of them was assayed by the loss of NcoI site (underlined) that covers the start ATG (bold) of *vll*. caCAGCTCAT<u>CC**ATG**G</u>tgcaGAGCATCTGATGGAGct *brca2* mutants were induced by CRISPR/Cas9 system. The sequence of gRNA targeting *brca2* is as follow. 5’TGGCTTGGATATGACACACC 3’ The *brca2* mutant allele, *brca2^sh515^*, contains an 83bp insertion at the amino acid 445 (exon 10) position introducing premature stop.

### Cell culture

MRC5 and U2OS cells were grown in Minimum Essential Media (MEM; Sigma-Aldrich) supplemented with 10% fetal bovine serum (Sigma-Aldrich), 2% L-glutamine (Sigma-Aldrich).

### Ionizing irradiation treatment in zebrafish

Zebrafish embryos were irradiated at 20 gray at 1dpf or 2 dpf with *γ*-ray using the irradiator (CIS bio international) at the Royal Hallamshire Hospital animal facility or with X-ray using the irradiator (Faxitron X-ray Corporation) at the Biomedical Science Department at the University of Sheffield.

### Injections into zebrafish embryos

CRISPR against *vhl* and *AR* was injected into the newly fertilized 1-cells stage embryos using microinjector (Pneumatic PicoPump WPI) at 100uM concentration at 1nl volume.

### Deep sequencing

The embryos from *vhl+/-* pair mating and *vhl+/-;vll-/-* pair mating were collected and the 15 embryos per each genotype were selected. After genomic DNAs were extracted from each group, the deep sequencing was performed with the primers shown below with three biological replicates. Forward primers for AR: TCGTCGGCAGCGTCAGATGTGTATAAGAGACAGCCT CCA AAG CAA AGG ACA CC (inner primer), ATGCCCCGTGATCTGAATGA (outer primer), Reverse primers for AR: GTCTCGTGGGCTCGGAGATGTGTATAAGAGACAGAAA TCAGTTGTGGCGCAGAG (inner primer), ACCCTGTGCACGGTGTATTA (outer primer), Forward primers for *vhl*: TCGTCGGCAGCGTCAGATGTGTATAAGAGACAG TGAAGCTTTAGTCTAACTCGGTG (inner primer), CACGAACCCACAAAAGTTGTTAT(outer primer), Reverse primers for *vhl*: GTCTCGTGGGCTCGGAGATGTGTATAAGAGACAG AATTCAGCATAATTTCACGAACC (inner primer), CTTCACCGACTCTCACAAGATTA (outer primer). Analysis of mutations frequencies and sizes was performed using BATCH-GE [49, 50]

### Immunostaining in zebrafish

Antibody staining was performed essentially according to the protocol. Embryos were fixed at -20°C with methanol:acetone (50%:50%) The embryos were permeabilised with 1% triton for 1 hour. The embryos were subsequently incubated in the blocking solution (2%Roche block, 5% calf serum, 1% DMSO). *γ*H2AX antibody (GTX127342, GeneTex,) was treated at 1:1000 dilution and incubated overnight. Secondary antibody(Alexa Flore 488 or Alexa Flore 546, Molecular Probe) was treated at 1:1000 and the embryos were incubated for 2 hours at room temperature. After the washing, the embryos were visualized using a confocal microscope.

### Chemical treatment

ATM inhibitor (KU-55933, Sigma-Aldrich) was treated both in zebrafish and cell cultures at 10uM concentration. HIF activators (FG 4592, Stratech or JNJ-42041935, Johnson and Johnson) were treated at 5uM and 100uM respectively in zebrafish and as indicated in cell cultures.

### TUNEL staining

3 hours after the irradiation with X-ray, the embryos were fixed with 4% PFA overnight at 4 °C. The embryos were washed 3 times for 5 minutes with PBS and permeabilised with 10 µg/ml proteinase K for 20 minutes. The fixed embryos were then washed and stained using the ApopTag Red in situ apoptosis detection kit (Chemicon).

### q-RCR for full length and short form of p53

The embryos from *vhl+/-* pair mating and *vhl+/-;vll-/-* pair mating were collected and raised to 2dpf. Then the embryos were irradiated with *γ*-ray at 20 gray dose. The embryos were pools according to their genotype and Total RNA was extracted. The specific TaqMan probes for full length and short form of p53 were designed and qPCR was performed after the first strand cDNA synthesis.

### Acridine Orange staining

Acridine Orange (A6014, Sigma Aldrich) was prepared at 1mg/ml (x100, stock solution) in miliQ water, and stored at -20 °C. The embryos were incubated in x1 Acridine Orange for 30 minutes in E3 fish medium. The embryos were washed three times each for 10 minutes.

### Immunostaining in the MRC5 and U2OS cells

70,000 of either MRC5 cells or U2OS cells were seeded onto the cover slips in the 24 well plates, and the cells were grown overnight at 37°C. Then the cells were treated with HIF activator for 3 hours followed by x-ray treatment either 5 or 2 gray dose. The irradiated cells were fixed with 4% PFA for 10 minutes after 30 minutes, 4 hours and 24 hours irradiation. The fixed cells were permeabilized with 0.2% tween and blocked for 1 hour with 3% BSA. The primary antibody gH2AX (mouse JWB301-Milipore) was added at 1:500 dilution and the cells were incubated in the primary antibody overnight at 4°C. Next day, anti-mouse Alexa Flour 594 (Invitrogen A11005) was added and the cells were incubated in the secondary antibody for one hour. The cells were counter stained with DAPI before mounting with mounting solution.

### Clonogenic studies

4000 cells per well were plated into the10cm plates. After leaving the cells to grow overnight at 37°C, cells were treated with chemicals, either ATMi at 10uM concentration alone or ATMi and HIF activators JNJ-42041935 at 100 uM or FG 4592 at 50uM and incubated at 37°C for 3 hours. Next CPT was added to the wells at 10nM or 25nM concentration, and cells were incubated 1 hour. The drugs were poured off and the fresh media were added to the plates. The cells were then returned to 37°C incubator for another 7 days for the colony formation. When the colonies are formed, the media were poured off and cells were fixed with 80% ethanol. After the cells were left to dry, they were stained with 1% methylene blue for 1 hour. Finally the plates were rinsed with water and colonies were counted.

### Statistical analysis

The data were analysed using the GraphPad program. All data were presented with the mean value of s.e.m. Statistical difference was analysed by the student’s t-test for two groups of data, one-way ANOVA for more than two groups. Two-way ANOVA test was performed to determine the effect of X-rays and mutations on the expression levels of p53 and *Δ*113p53. The significance of Hif activation on the survival of zebrafish embryos was tested by chi-square test. *p<0.05* was considered statistically significant.

### Imaging

The embryos were visualized using Leica dissecting microscope and the images were captured using Leica Application System v4.9.0 software. The confocal images were acquired using Olympus FV1000 available at the light microscope facility at the Biomedical Science Department at the University of Sheffield.

## Abbreviations

VHL: Von Hippel Lindau
ccRCC: clear cell renal cell carcinoma
HIF: Hypoxia Inducible Factor
PHD: Prolyl Hydroxylase
HRE: HIF responsive element
DSB: double strand break
HR: homologous recombination
A-T: Ataxia -Telangiectasia
MMR: mismatch repair
LOH: loss of heterozygosity
TUNEL: terminal deoxynucleotidyl transferase dUTP nick end labelling
CPT: Camptothecin
CNS: central nervous system
ATM: ATM serine/threonine kinase
ATMi: ATM inhibitor
mpi: minutes post irradiation
APTX: aprataxin
ATLD: A-T like disease
SCAN1: spinocerebellar ataxia with axonal neuropathy
ZFN: zinc finger nucleases

## Author Contributions

Kim HR, wrote the paper, designed and executed experiments.

Santhakumar K, wrote the paper, designed and executed experiments.

Markham E, executed experiments.

Baldera D, executed experiments.

Bryant HE, designed experiments and commented on manuscript.

El-Khamisy SF, designed experiments and commented on manuscript.

van Eeden FJ, designed experiments, executed experiments, wrote the manuscript.

## Acknowledgements

We thank Dan Harris and Shih-Chieh Chiang for help with cell culture work. The Bateson Centre aquaria staff kindly provided care for zebrafish. We also would like to thank Elisabeth Kugler for help with *γ*H2AX foci counting script.

## Bibliography

1. Gossage L, Eisen T, Maher ER. VHL, the story of a tumour suppressor gene. Nat Rev Cancer. 2015; 15: 55–64. doi: 10.1038/nrc3844.

2. Maher ER, Neumann HP, Richard S. von Hippel-Lindau disease: a clinical and scientific review. Eur J Hum Genet. 2011; 19: 617–23. doi: 10.1038/ejhg.2010.175.

3. Jaakkola P, Mole DR, Tian YM, Wilson MI, Gielbert J, Gaskell SJ, von Kriegsheim A, Hebestreit HF, Mukherji M, Schofield CJ, Maxwell PH, Pugh CW, Ratcliffe PJ. Targeting of HIF-alpha to the von Hippel-Lindau ubiquitylation complex by O2-regulated prolyl hydroxylation. Science. 2001; 292: 468–72. doi: 10.1126/science.1059796.

4. Kaelin WG, Jr. The VHL Tumor Suppressor Gene: Insights into Oxygen Sensing and Cancer. Trans Am Clin Climatol Assoc. 2017; 128: 298–307. doi:

5. Semenza GL. Hypoxia-inducible factors in physiology and medicine. Cell. 2012; 148: 399–408. doi: 10.1016/j.cell.2012.01.021.

6. Li M, Kim WY. Two sides to every story: the HIF-dependent and HIF-independent functions of pVHL. J Cell Mol Med. 2011; 15: 187–95. doi: 10.1111/j.1582-4934.2010.01238.x.

7. Metcalf JL, Bradshaw PS, Komosa M, Greer SN, Stephen Meyn M, Ohh M. K63-ubiquitylation of VHL by SOCS1 mediates DNA double-strand break repair. Oncogene. 2014; 33: 1055–65. doi: 10.1038/onc.2013.22.

8. Scanlon SE, Hegan DC, Sulkowski PL, Glazer PM. Suppression of homology-dependent DNA double-strand break repair induces PARP inhibitor sensitivity in VHL-deficient human renal cell carcinoma. Oncotarget. 2018; 9: 4647–60. doi: 10.18632/oncotarget.23470.

9. Hoffman MA, Ohh M, Yang H, Klco JM, Ivan M, Kaelin WG, Jr. von Hippel-Lindau protein mutants linked to type 2C VHL disease preserve the ability to downregulate HIF. Hum Mol Genet. 2001; 10: 1019–27. doi:

10. Percy MJ, Furlow PW, Lucas GS, Li X, Lappin TR, McMullin MF, Lee FS. A gain-of-function mutation in the HIF2A gene in familial erythrocytosis. N Engl J Med. 2008; 358: 162–8. doi: 10.1056/NEJMoa073123.

11. Ang SO, Chen H, Hirota K, Gordeuk VR, Jelinek J, Guan Y, Liu E, Sergueeva AI, Miasnikova GY, Mole D, Maxwell PH, Stockton DW, Semenza GL, et al. Disruption of oxygen homeostasis underlies congenital Chuvash polycythemia. Nat Genet. 2002; 32: 614–21. doi: 10.1038/ng1019.

12. Lopez-Otin C, Blasco MA, Partridge L, Serrano M, Kroemer G. The hallmarks of aging. Cell. 2013; 153: 1194–217. doi: 10.1016/j.cell.2013.05.039.

13. Menck CF, Munford V. DNA repair diseases: What do they tell us about cancer and aging? Genet Mol Biol. 2014; 37: 220–33. doi:

14. Spry M, Scott T, Pierce H, D’Orazio JA. DNA repair pathways and hereditary cancer susceptibility syndromes. Front Biosci. 2007; 12: 4191–207. doi:

15. Gatti R, Perlman S. (1993). Ataxia-Telangiectasia. In: Adam MP, Ardinger HH, Pagon RA, Wallace SE, Bean LJH, Stephens K and Amemiya A, eds. GeneReviews((R)). (Seattle (WA).

16. Bryant HE, Schultz N, Thomas HD, Parker KM, Flower D, Lopez E, Kyle S, Meuth M, Curtin NJ, Helleday T. Specific killing of BRCA2-deficient tumours with inhibitors of poly(ADP-ribose) polymerase. Nature. 2005; 434: 913–7. doi: 10.1038/nature03443.

17. Borrell B. How accurate are cancer cell lines? Nature. 2010; 463: 858. doi: 10.1038/463858a.

18. Wilding JL, Bodmer WF. Cancer cell lines for drug discovery and development. Cancer Res. 2014; 74: 2377–84. doi: 10.1158/0008-5472.CAN-13-2971.

19. Xia Y, Jiang L, Zhong T. The role of HIF-1alpha in chemo-/radioresistant tumors. Onco Targets Ther. 2018; 11: 3003–11. doi: 10.2147/OTT.S158206.

20. Calvo-Asensio I, Dillon ET, Lowndes NF, Ceredig R. The Transcription Factor Hif-1 Enhances the Radio-Resistance of Mouse MSCs. Front Physiol. 2018; 9: 439. doi: 10.3389/fphys.2018.00439.

21. Lu C, El-Deiry WS. Targeting p53 for enhanced radio- and chemo-sensitivity. Apoptosis. 2009; 14: 597–606. doi: 10.1007/s10495-009-0330-1.

22. Howe K, Clark MD, Torroja CF, Torrance J, Berthelot C, Muffato M, Collins JE, Humphray S, McLaren K, Matthews L, McLaren S, Sealy I, Caccamo M, et al. The zebrafish reference genome sequence and its relationship to the human genome. Nature. 2013; 496: 498–503. doi: 10.1038/nature12111.

23. van Rooijen E, Voest EE, Logister I, Korving J, Schwerte T, Schulte-Merker S, Giles RH, van Eeden FJ. Zebrafish mutants in the von Hippel-Lindau tumor suppressor display a hypoxic response and recapitulate key aspects of Chuvash polycythemia. Blood. 2009; 113: 6449–60. doi: 10.1182/blood-2008-07-167890.

24. van Rooijen E, Voest EE, Logister I, Bussmann J, Korving J, van Eeden FJ, Giles RH, Schulte-Merker S. von Hippel-Lindau tumor suppressor mutants faithfully model pathological hypoxia-driven angiogenesis and vascular retinopathies in zebrafish. Dis Model Mech. 2010; 3: 343–53. doi: 10.1242/dmm.004036.

25. Kim HR, Greenald D, Vettori A, Markham E, Santhakumar K, Argenton F, van Eeden F. Zebrafish as a model for von Hippel Lindau and hypoxia-inducible factor signaling. Methods Cell Biol. 2017; 138: 497–523. doi: 10.1016/bs.mcb.2016.07.001.

26. Santhakumar K, Judson EC, Elks PM, McKee S, Elworthy S, van Rooijen E, Walmsley SS, Renshaw SA, Cross SS, van Eeden FJ. A zebrafish model to study and therapeutically manipulate hypoxia signaling in tumorigenesis. Cancer Res. 2012; 72: 4017–27. doi: 10.1158/0008-5472.CAN-11-3148.

27. Greenald D, Jeyakani J, Pelster B, Sealy I, Mathavan S, van Eeden FJ. Genome-wide mapping of Hif-1alpha binding sites in zebrafish. BMC Genomics. 2015; 16: 923. doi: 10.1186/s12864-015-2169-x.

28. Pescador N, Cuevas Y, Naranjo S, Alcaide M, Villar D, Landazuri MO, Del Peso L. Identification of a functional hypoxia-responsive element that regulates the expression of the egl nine homologue 3 (egln3/phd3) gene. Biochem J. 2005; 390: 189–97. doi: 10.1042/BJ20042121.

29. Pena-Llopis S, Christie A, Xie XJ, Brugarolas J. Cooperation and antagonism among cancer genes: the renal cancer paradigm. Cancer Res. 2013; 73: 4173–9. doi: 10.1158/0008-5472.CAN-13-0360.

30. Ryan AJ, Squires S, Strutt HL, Johnson RT. Camptothecin cytotoxicity in mammalian cells is associated with the induction of persistent double strand breaks in replicating DNA. Nucleic Acids Res. 1991; 19: 3295–300. doi:

31. Ashour ME, Atteya R, El-Khamisy SF. Topoisomerase-mediated chromosomal break repair: an emerging player in many games. Nat Rev Cancer. 2015; 15: 137–51. doi: 10.1038/nrc3892.

32. Liao C, Beveridge R, Hudson JJR, Parker JD, Chiang SC, Ray S, Ashour ME, Sudbery I, Dickman MJ, El-Khamisy SF. UCHL3 Regulates Topoisomerase-Induced Chromosomal Break Repair by Controlling TDP1 Proteostasis. Cell Rep. 2018; 23: 3352–65. doi: 10.1016/j.celrep.2018.05.033.

33. Roe JS, Youn HD. The positive regulation of p53 by the tumor suppressor VHL. Cell Cycle. 2006; 5: 2054–6. doi: 10.4161/cc.5.18.3247.

34. Gong L, Gong H, Pan X, Chang C, Ou Z, Ye S, Yin L, Yang L, Tao T, Zhang Z, Liu C, Lane DP, Peng J, et al. p53 isoform Delta113p53/Delta133p53 promotes DNA double-strand break repair to protect cell from death and senescence in response to DNA damage. Cell Res. 2015; 25: 351–69. doi: 10.1038/cr.2015.22.

35. Rodriguez-Mari A, Wilson C, Titus TA, Canestro C, BreMiller RA, Yan YL, Nanda I, Johnston A, Kanki JP, Gray EM, He X, Spitsbergen J, Schindler D, et al. Roles of brca2 (fancd1) in oocyte nuclear architecture, gametogenesis, gonad tumors, and genome stability in zebrafish. PLoS Genet. 2011; 7: e1001357. doi: 10.1371/journal.pgen.1001357.

36. Shiloh Y, Ziv Y. The ATM protein kinase: regulating the cellular response to genotoxic stress, and more. Nat Rev Mol Cell Biol. 2013; 14: 197–210. doi:

37. Smith K, Gunaratnam L, Morley M, Franovic A, Mekhail K, Lee S. Silencing of epidermal growth factor receptor suppresses hypoxia-inducible factor-2-driven VHL-/- renal cancer. Cancer Res. 2005; 65: 5221–30. doi: 10.1158/0008-5472.CAN-05-0169.

38. Hughes MD, Kapllani E, Alexander AE, Burk RD, Schoenfeld AR. HIF-2alpha downregulation in the absence of functional VHL is not sufficient for renal cell differentiation. Cancer Cell Int. 2007; 7: 13. doi: 10.1186/1475-2867-7-13.

39. Nyhan MJ, O’Sullivan GC, McKenna SL. Role of the VHL (von Hippel-Lindau) gene in renal cancer: a multifunctional tumour suppressor. Biochem Soc Trans. 2008; 36: 472–8. doi: 10.1042/BST0360472.

40. Loeb LA. A mutator phenotype in cancer. Cancer Res. 2001; 61: 3230–9. doi:

41. Loeb LA, Bielas JH, Beckman RA. Cancers exhibit a mutator phenotype: clinical implications. Cancer Res. 2008; 68: 3551–7; discussion 7. doi: 10.1158/0008-5472.CAN-07-5835.

42. Biton S, Barzilai A, Shiloh Y. The neurological phenotype of ataxia-telangiectasia: solving a persistent puzzle. DNA Repair (Amst). 2008; 7: 1028–38. doi: 10.1016/j.dnarep.2008.03.006.

43. Stewart GS, Maser RS, Stankovic T, Bressan DA, Kaplan MI, Jaspers NG, Raams A, Byrd PJ, Petrini JH, Taylor AM. The DNA double-strand break repair gene hMRE11 is mutated in individuals with an ataxia-telangiectasia-like disorder. Cell. 1999; 99: 577–87. doi:

44. Takashima H, Boerkoel CF, John J, Saifi GM, Salih MA, Armstrong D, Mao Y, Quiocho FA, Roa BB, Nakagawa M, Stockton DW, Lupski JR. Mutation of TDP1, encoding a topoisomerase I-dependent DNA damage repair enzyme, in spinocerebellar ataxia with axonal neuropathy. Nat Genet. 2002; 32: 267–72. doi: 10.1038/ng987.

45. Date H, Onodera O, Tanaka H, Iwabuchi K, Uekawa K, Igarashi S, Koike R, Hiroi T, Yuasa T, Awaya Y, Sakai T, Takahashi T, Nagatomo H, et al. Early-onset ataxia with ocular motor apraxia and hypoalbuminemia is caused by mutations in a new HIT superfamily gene. Nat Genet. 2001; 29: 184–8. doi: 10.1038/ng1001-184.

46. Moreira MC, Barbot C, Tachi N, Kozuka N, Uchida E, Gibson T, Mendonca P, Costa M, Barros J, Yanagisawa T, Watanabe M, Ikeda Y, Aoki M, et al. The gene mutated in ataxia-ocular apraxia 1 encodes the new HIT/Zn-finger protein aprataxin. Nat Genet. 2001; 29: 189–93. doi: 10.1038/ng1001-189.

47. Besarab A, Chernyavskaya E, Motylev I, Shutov E, Kumbar LM, Gurevich K, Chan DT, Leong R, Poole L, Zhong M, Saikali KG, Franco M, Hemmerich S, et al. Roxadustat (FG-4592): Correction of Anemia in Incident Dialysis Patients. J Am Soc Nephrol. 2016; 27: 1225–33. doi: 10.1681/ASN.2015030241.

48. Provenzano R, Besarab A, Sun CH, Diamond SA, Durham JH, Cangiano JL, Aiello JR, Novak JE, Lee T, Leong R, Roberts BK, Saikali KG, Hemmerich S, et al. Oral Hypoxia-Inducible Factor Prolyl Hydroxylase Inhibitor Roxadustat (FG-4592) for the Treatment of Anemia in Patients with CKD. Clin J Am Soc Nephrol. 2016; 11: 982–91. doi: 10.2215/CJN.06890615.

49. Boel A, Steyaert W, De Rocker N, Menten B, Callewaert B, De Paepe A, Coucke P, Willaert A. BATCH-GE: Batch analysis of Next-Generation Sequencing data for genome editing assessment. Sci Rep. 2016; 6: 30330. doi: 10.1038/srep30330.

50. Boel A, Steyaert W, De Rocker N, Menten B, Callewaert B, De Paepe A, Coucke P, Willaert A. Publisher Correction: BATCH-GE: Batch analysis of Next-Generation Sequencing data for genome editing assessment. Sci Rep. 2018; 8: 15845. doi: 10.1038/s41598-018-33869-y.

