## Supplementary data for "The investigation of the role of VHL-HIF signaling in DNA repair and apoptosis in zebrafish"

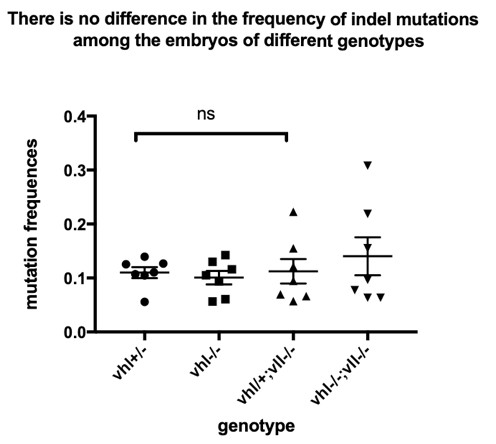

**Supplementary Fig. 1** **There was no difference in the frequency of small indel mutations, created by CRISPR injection against *AR*, among *vhl-/-,* *vhl-/-;vll-/-* embryos and their siblings.** Embryos were collected from *vhl+/-* and *vhl+/-;vll-/-* pair mating and were injected with CRISPR targeting *vhl* and *AR*. The genomic DNA around the target sites was amplified by PCR and sequenced by deep sequencing. This revealed that there was no significant difference in small indel mutation frequency between embryos with different genotypes. ^ns^ *p>0.05*, one way ANOVA.

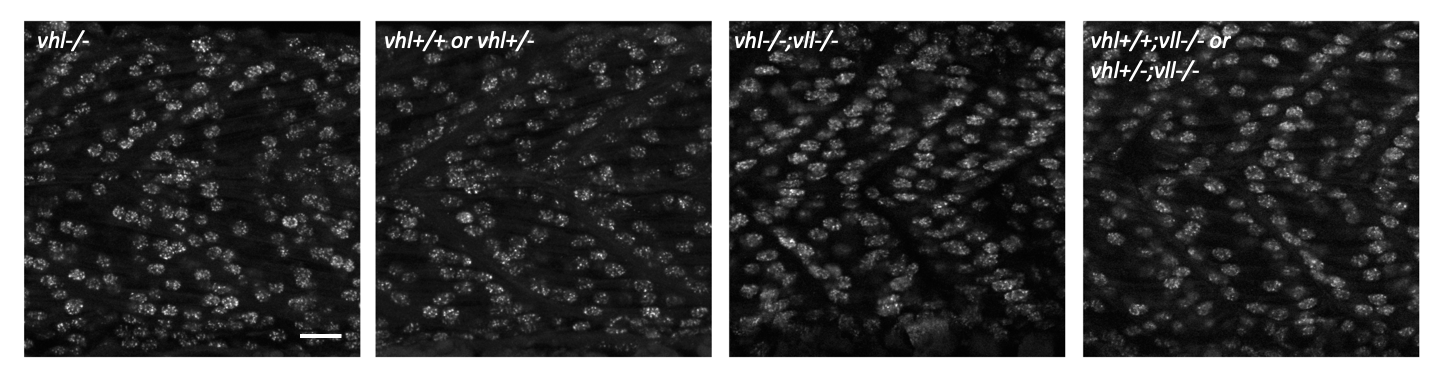

**Supplementary fig. 2**  **γH2AX foci were equally formed in *vhl-/-*, *vhl-/-;vll-/-* embryos and their siblings immediately after the irradiation.** The embryos from *vhl+/-* and *vhl+/-;vll-/-* incrosses were collected and irradiated with X-ray at 24hpf. Immediately after the irradiation the embryos were fixed and the γH2AX foci formation was examined. It revealed that all embryos of different genotypes showed equally distributed γH2AX foci formation, indicating the DNA damage is equally introduced in these embryos. Scale bar: 20μm

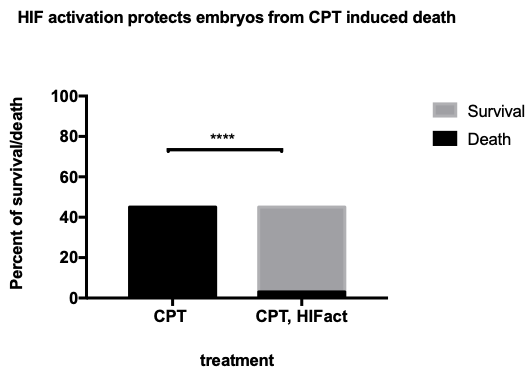

**Supplementary Fig. 3 Hif activator treated embryos were protected from CPT induced death.** When the wild type embryos were treated with 20nM CPT overnight, all embryos died by 5dpf. In contrast, the majority of embryos survived on 5dpf when Hif activator was treated with CPT (see Fig. 5 E and F). ^****^*p<0.0001*, chi-square test.
